# Astroglial TNFR2 signaling regulates hippocampal synaptic function and plasticity in a sex dependent manner

**DOI:** 10.1101/2025.03.13.643110

**Authors:** Brianna N. Carney, Placido Illiano, Taylor M. Pohl, Haritha L. Desu, Shwetha Mudalegundi, Andoni I. Asencor, Shika Jwala, Maureen C. Ascona, Praveen K. Singh, David J. Titus, Burcu A. Pazarlar, Lei Wang, Laura Bianchi, Jens D. Mikkelsen, Coleen M. Atkins, Kate L. Lambertsen, Roberta Brambilla

**Author notes:** Correspondence, The Miami Project to Cure Paralysis, Department of Neurological Surgery University of Miami Miller School of Medicine, 1095 NW 14^th^ Terrace, Miami, Florida 33136, USA.

## Abstract

Astrocytes participate in synaptic transmission and plasticity through tightly regulated, bidirectional communication with pre- and post-synaptic neurons, as well as microglia and oligodendrocytes. A key component of astrocyte-mediated synaptic regulation is the cytokine tumor necrosis factor (TNF). TNF signals via two cognate receptors, TNFR1 and TNFR2, both expressed in astrocytes. While TNFR1 signaling in astrocytes has been long demonstrated to be necessary for physiological synaptic function, the role of astroglial TNFR2 has never been explored. Here, we demonstrate that astroglial TNFR2 is essential for maintaining hippocampal synaptic function and plasticity in physiological conditions. Indeed, *Gfap^creERT2^:Tnfrsf1b^fl/fl^* mice with selective ablation of TNFR2 in astrocytes exhibited dysregulated expression of neuronal and glial proteins (e.g., SNARE complex molecules, glutamate receptor subunits, glutamate transporters) essential for hippocampal synaptic transmission and plasticity. Hippocampal astrocytes sorted from *Gfap^creERT2^:Tnfrsf1b^fl/fl^* mice displayed downregulation of genes and pathways implicated in synaptic plasticity, as well as astrocyte-neuron and astrocyte-oligodendrocyte communication. These alterations were accompanied by increased glial reactivity and impaired astrocyte calcium dynamics, and ultimately translated into functional deficits, specifically impaired long-term potentiation (LTP) and cognitive functions. Notably, male *Gfap^creERT2^:Tnfrsf1b^fl/fl^* mice exhibited more pronounced hippocampal synaptic and cellular alterations, suggesting sex-dependent differences in astroglial TNFR2 regulation of synaptic function. Together, these findings indicate that TNFR2 signaling in astrocytes is essential for proper astrocyte-neuron communication at the basis of synaptic function, and that this is regulated in a sex-dependent manner.

## 1. Introduction

Synaptic transmission and plasticity, the latter defined as long-lasting activity-dependent changes in the efficacy of synaptic transmission [1], rely on the integrated and bidirectional communication between neurons and glia. While neurons form the central components of this circuitry, glia, and particularly astrocytes, have essential functions [2]. Astrocytes are integral parts of the synapse [3, 4]. Their complex processes are tightly connected to pre- and post-synaptic terminals enabling the fine-tuning of neuronal synaptic transmission [5–7]. This occurs via the regulation of gliotransmitter release (e.g. glutamate, GABA, ATP, D-serine) [8], glutamate buffering [9], and synaptic formation and elimination [10–13]. Hence, when astrocyte-neuron communication is disturbed, synaptic transmission and plasticity are impaired, leading to compromised neural functions. This includes alterations in cognitive processes, whose neurobiological basis is indeed the plastic adaptation of synaptic transmission, particularly within the hippocampus.

A key modulator of synaptic transmission and plasticity is the cytokine tumor necrosis factor (TNF) [14]. TNF exists in two forms: a native transmembrane form, tmTNF, and a soluble form, solTNF. tmTNF functions via cell-to-cell contact and, when cleaved by the metalloproteinase ADAM17, sheds solTNF [15, 16]. TNF binds to two receptors, TNFR1 and TNFR2, which differ in ligand affinity, downstream signaling and cellular expression. solTNF preferentially binds to and activates TNFR1; tmTNF binds to both receptors but can only activate TNFR2 downstream signaling [17, 18]. TNFR1 and TNFR2 are expressed in astrocytes [19–21], both in physiological and pathological conditions [22, 23]. TNF-mediated signaling in astrocytes is a major contributor to astrocyte-dependent modulation of synaptic transmission/plasticity in health and disease [22, 24]. Under physiological TNF levels, astroglial TNF signaling through TNFR1 is key to regulating homeostatic synaptic plasticity such as synaptic scaling [25], glutamate release [26, 27], and glutamate and GABA receptor trafficking [28, 29]. Under neuroinflammatory conditions where TNF levels are elevated, overactivated astroglial TNFR1 signaling triggers astrocyte-to-neuron cascades that cause persistent functional modifications of hippocampal excitatory synapses, leading to impairment of long-term potentiation (LTP) and consequent deficits in learning and memory [30].

Remarkably, astrocytic TNFR2 signaling has never been studied for its role in synaptic function despite being expressed in astrocytes, and being attributed a protective and reparative function when expressed in microglia and oligodendroglia, both in health and neurological disease [19, 21, 31, 32]. On this basis, we set out to address the role of astroglial TNFR2 postulating its participation in sustaining proper physiologic synaptic transmission and plasticity. To this end, we employed *Gfap^creERT2^:Tnfrsf1b^fl/fl^* transgenic mice with astrocyte-specific ablation of TNFR2 where we observed, in naïve conditions, functional alterations ranging from impaired hippocampal synaptic plasticity to cognitive deficits. These were paralleled by aberrant expression of hippocampal molecules implicated in synaptic function, as well as phenotypic changes in astroglia and microglia associated with increased reactivity. Notably, we observed more pronounced effects in male mice, indicating their stronger reliance on TNFR2 for homeostatic regulation. Together, these data demonstrate the essential contribution of astroglial TNFR2 to hippocampal synaptic function and underscore the crucial role of astroglial TNF signaling as a whole, both via TNFR2 and TNFR1, as a regulator of physiological synaptic transmission, plasticity, and cognition.

## 2. Materials and methods

### 2.1. Mice

Mice with astrocyte specific conditional ablation of the *Tnfrsf1b* gene (encoding for the TNFR2 protein) were generated by crossing *Gfap^creERT2^* mice [33], kindly provided by Dr. Ken McCarthy, with *Tnfrsf1b^fl/fl^* mice. *Tnfrsf1b^fl/fl^* mice were generated in our laboratory as previously described [34]. In all experiments, *Tnfrsf1b^fl/fl^* littermates were used as controls. Genotyping of the *Gfap^creERT2^* transgenic allele was conducted with the following primers: forward = 5’ ggtcgatgcaacgagtgatgagg 3’; reverse = 5’ gctaagtgccttctctacacctgcg 3’. Genotyping of the *Tnfrsf1b^fl^* transgenic allele was conducted with the following primers: forward = 5′ ttgggtctagaggtggcgcagc 3′; reverse = 5′ ggccaggaagtgggttactttagggc 3′. Recombination was induced by intraperitoneal (i.p.) injections of tamoxifen (125 mg/kg) for 5 consecutive days. This was followed by a 3-week wash out period before starting the experiments. Recombination efficiency was tested in *Gfap^creERT2^:Eyfp:Tnfrsf1b^fl/fl^* mice, obtained from crossing *Gfap^creERT2^:Tnfrsf1b^fl/fl^* mice with *Rosa26-stop-Eyfp* reporter mice (Jackson Laboratories, stock # 006148). Adult (2-4 months) male and female mice were used in all *in vivo* studies. C57BL/6J wild type (WT) and germline *Tnfrsf1b*^-/-^ (Jackson Laboratories, stock # 002620) pups (postnatal day 3-5) were used to generate primary astrocyte cultures for *in vitro* studies.

All mice were group-housed (maximum 5 mice/cage) in the Animal Core Facility of The Miami Project to Cure Paralysis in a virus/antigen-free, temperature- and humidity-controlled room with a 12 hr light/dark cycle. Mice were provided with water and food *ad libitum*. All experiments were performed according to protocols and guidelines approved by the Institutional Animal Care and Use Committee of the University of Miami.

### 2.2. Behavioral assessments

#### Open field test

Exploratory behavior and locomotion were assessed with the open field test in adult mice as previously described [34]. Mice were placed in an odor-free, non-transparent squared arena [dimensions in cm: 72 (width), 72 (depth), 40 (height)] for three consecutive days and for periods of 5, 10 and 10 min, respectively. The first and second days in the arena were considered habituation sessions, and the third day was the test day. Mice were habituated to the room 1 hr prior to testing. The arena was divided into four zones (wall, outer, intermediate, and center), and mouse behavior during exploration was recorded using a high-resolution color video camera (SSC-DC378P, Biosite, Stockholm, Sweden). The investigator left the room during testing to minimize factors that might affect mouse performance (e.g., noise, smell). Tests were conducted at the same time each day and by the same investigator. Distance traveled, velocity and zone transitions were analyzed using the EthoVision XT 11.5 Video Tracking Software (Noldus Information Technology, Leesburg, VA, USA) [35]. The experimenter was blinded to genotype and experimental conditions.

#### Rotarod test

Locomotor coordination was assessed in adult mice with the LE8200 rotarod apparatus (Panlab) as previously described [34]. Mice underwent pre-training for 5 consecutive days, followed by one rest day, and the test day (day 7). Prior to each session, mice were habituated to the room for 1 hr. Exclusion criteria were established in that mice unable to maintain their balance for a minimum of 60 sec on the rotating rod set at a constant speed of 4 rpm were excluded from the study. For the experiment (both pre-training and test days), the rod was set to rotate at increasing speed and constant acceleration from 4 to 40 rpm over a 10 min period. On test day, each mouse performed a total of four trials. After each trial, mice were transferred back to their home cage for at least 20 min before the next trial to minimize exhaustion and stress. For each trial, the total time spent on the rod was recorded by an experimenter blinded to genotype and experimental conditions. The final outcome was calculated as an average of the four trials.

#### Novel object recognition test

Long-term memory was assessed in adult mice with the novel object recognition test as previously described [36], with minor modifications. Mice were placed in an odor-free, non-transparent rectangular arena [dimensions in cm: 37.5 (width), 26 (depth), 20 (height)] for three consecutive days: on day 1 (habituation), mice were placed in the empty arena for 5 min to familiarize with the environment; on day 2 (exploration), mice were placed in the arena fitted with two identical round-shaped objects equally spaced from the walls for a total of 10 min; on day 3 (test), mice were re-introduced into the arena where one of the round-shaped objects was replaced with a rectangular-shaped novel object for a total of 10 min. Objects were equally spaced from the walls, and the position of the novel object (left v*s* right) was randomly assigned to each genotype and condition. Mice were habituated to the room 1 hr prior to testing. The investigator left the room during testing to minimize factors that might affect mouse performance (e.g., noise, smell). The test was performed at the same time each day and by the same investigator. Activity in the arena was recorded using a high-resolution color video camera (SSC-DC378P, Biosite, Stockholm, Sweden) and analyzed with the EthoVision XT 11.5 Video Tracking Software (Noldus Information Technology, Leesburg, VA, USA) [35]. Total time spent exploring each object was recorded, and the discrimination index calculated (Time with novel object – Time with familiar object / Total time).

#### Fear conditioning test

Associative memory was assessed in adult mice with the fear conditioning test as previously described [37]. Mice were placed in an isolation cubicle (Coulbourn Instruments) containing a fear conditioning chamber (Coulbourn Instruments) connected to an electric grid floor for three consecutive days: on day 1 (habituation), mice were placed inside the fear conditioning apparatus for 10 min to familiarize with the environment; on day 2 (training), mice were returned to the apparatus for a 210 sec long training session. At 120 sec, mice were exposed to an auditory cue lasting 30 sec (85 dB, 2kHz), followed by a 0.7 mA foot shock during the last 3 sec of the auditory cue; on day 3 (test), mice were placed in the apparatus for 5 min to measure the freezing response resulting from context-based associative memory. Mice were then removed from the apparatus for 1 h, during which the chamber environment was modified by replacing walls and grid floor, and by introducing a novel scent. Mice were placed back into the apparatus and their freezing response was measured for 3 min before, and 3 min during exposure to the auditory cue to test for cue-based associative memory. Freezing time was quantified with the video-based analysis software FreezeFrame 5 (Actimetrics) and expressed as percentage of total time. The percentage of time spent freezing in the first 90 sec of the training day (day 1) was used as baseline freezing percentage. The test was performed at the same time each day and by the same investigator, blind to the genotype.

#### Morris water maze

Spatial memory was assessed in adult mice with the Morris water maze test as previously described [38]. Mice were trained for four consecutive days (acquisition) in a pool containing a hidden platform located just below the water surface, and with spatial cues placed by each of the four pool quadrants. Mice were introduced into the pool at different starting points and allowed to swim until they reached the hidden platform. Each mouse underwent four 65 sec trials per day (each at a different starting point), with 25 min rest time between trials. The average time to reach the hidden platform during the acquisition days was quantified as a measure of spatial learning. On day 5 (probe trial), the hidden platform was removed, and the time to reach the target area was quantified as a measure of spatial memory. Time to platform and time to target area were extrapolated using the EthoVision XT 11.5 Video Tracking Software (Noldus Information Technology, Leesburg, VA, USA) [35]. Mice were habituated to the room 1 hr prior to testing, and the test was performed at the same time each day by the same investigator, who was blind to the genotype.

#### Light-dark transition test

Anxiety associated behavior was assessed with the light-dark transition test in adult mice as previously described [39], with minor modifications. In a completely dark room, mice were placed in a testing apparatus [dimensions, in cm: 37.5 (width), 26 (depth), 20 (height)] consisting of two separate chambers connected through a small opening closed by a removable door. One chamber was brightly illuminated with white walls, the other dimly lit with black walls. At the start of the test, with the connecting door shut, mice were placed in the dark chamber for 1 min. The door was then opened and mice allowed to freely explore both chambers for 10 min. Activity in the illuminated chamber was recorded with a high-resolution color video camera (SSC-DC378P, Biosite, Stockholm, Sweden). Total distance traveled (m), velocity (m/s), total movement time (sec), number of transitions between chambers, time spent in the light chamber (sec), latency to light chamber exploration (sec), number of wall explorations, and wall exploration time (sec) were estimated using the EthoVision XT 11.5 Video Tracking Software (Noldus Information Technology) [35]. The investigator left the room during testing to minimize factors that might affect mouse performance. The test was performed at the same time each day and by the same investigator blind to genotype.

### 2.3. Immunohistochemistry and stereological analysis

After transcardial perfusion with 0.1 M PBS followed by 4% paraformaldehyde (PFA, made in 0.1 M PBS), spinal cords and brains were dissected and post-fixed for 3 h in 4% PFA. Tissues were then cryoprotected in 0.1 M PBS + 30% sucrose and cryostat-cut into 30 μm-thick serial sections. Tissue sections were blocked with 5% normal goat serum (NGS) in 0.1 M PBS + 0.4% Triton-X-100, then incubated overnight at 4°C with antibodies against green fluorescent protein (GFP) (chicken, 1:1000, Abcam, #ab13970), GFAP (rat, 1:500, Invitrogen, #13-0300), Iba1 (rabbit, 1:500, WAKO, #019-19741), or NeuN (guinea pig, 1:200, Synaptic Systems, #266-006). Immunoreactivity was visualized with AlexaFluor 488, 594, and 647 species-specific secondary antibodies (1:750, Invitrogen). Sections were stained with DAPI (1:10,000, Invitrogen), rinsed, coverslipped with Flouromount mounting medium (Sigma), and imaged on a Zeiss LSM 800 confocal or BZ-X800 Keyence fluorescence microscope. For unbiased quantification of hippocampal astrocytes, microglia and neurons, stereological analysis was employed. For each animal, four 30 μm-thick serial sections taken at 300 μm intervals were imaged and analyzed with an Olympus BX51 fluorescence microscope at a 63X magnification using StereoInvestigator® software (MicroBrightfield Inc.), as previously published [40, 41]. The total number of cells per mm^3^ of tissue was estimated using the Optical Fractionator function which allows for systematic random sampling. The investigator was blinded to the genotype.

### 2.4. Sholl analysis

Sholl analysis was performed on NeuN^+^, Iba1^+^, and GFAP^+^ cells in the CA1 region of the hippocampus. Following immunostaining of hippocampal tissues (see paragraph 2.3), sections were imaged at 40x or 100x magnification on a BZ-X800 Keyence fluorescence microscope. Six cells/group were randomly selected for morphological analysis, ensuring that cell processes did not overlap with neighboring cells. Manual tracing of cell processes was implemented with ImageJ software by an investigator blinded to the genotype. The number of process intersections was calculated at an increasing distance from the cell nucleus (30 μm increments for NeuN^+^ cells, 4 μm increments for GFAP^+^ and Iba1^+^ cells). Total process length/cell, average process length/cell, total number of processes/cell, and maximum intersection radius/cell were also quantified.

### 2.5. Autoradiography of [^3^H]PBR28 binding for translocator protein (TSPO) detection

Glial reactivity was evaluated through autoradiographic quantification of the [^3^H]PBR28 radioligand bound to TSPO in brain sections as previously described [42, 43], with minor modifications. PBS-perfused brains were dissected out, flash frozen in liquid nitrogen, and cryostat-cut into 30 μm-thick serial sections. Tissue sections were washed twice for 10 min in pre-incubation buffer containing 50 mM Tris-base and 0.5% bovine serum albumin (BSA), then incubated for 1 h at RT in incubation buffer consisting of 5 mM Tris-HCl with 0.5% BSA, 5 mM MgCl_2_, 2 mM EGTA, and 3 nM [^3^H]PBR28 (Tritec AG, Switzerland; specific activity: 80.7 Ci/mmol; concentration: 1 mCi/ml). Slides were then rinsed with ice cold pre-incubation buffer, dipped in ice cold distilled water, fixed overnight in a PFA vapor chamber at 4°C, and dried in a silica gel desiccator for 45 min. Sections were exposed to a tritium-sensitive Fuji IP BAS-TR 2040 imaging plate (Fujifilm) for 48 hr at 4°C, along with tritium standards ([^3^H] microscale ART0123 (0-489.1 nCi/mg) and ART0123B (3-109.4 nCi/mg), American Radiolabeled Chemicals) in a radiation-shielded cassette (Fujifilm). The imaging plate was then scanned with a Typhoon IP Biomolecular Imager (GE Healthcare) at a pixel size of 25 μm. To determine the specific radioligand binding to the target, in an adjacent section displacement of [^3^H]PBR28 was performed by exposure to 10 μM PK11195 (provided by Lundbeck A/S, Batch number A0031BF) dissolved in incubation buffer containing 3 nM [^3^H]PBR28. Binding was quantified by measuring the mean optical density (OD) in each region of interest (ROI). Using ImageJ Rodbard software, the Grey radioactivity values from tritium standards were used to convert OD values from ROIs into Gray radioactivity values. The decay-corrected specific activity of the radioligand was used to convert radioactivity (nCi/mg) to binding values in tissue (fmol/mg).

### 2.6. Multiplex analysis

Multiplex analysis was performed in hippocampal tissue as previously described [41]. After transcardial perfusion with 0.1 M PBS, the hippocampus was dissected and homogenized in 300 μl of RIPA buffer containing phosphatase and protease inhibitors (Sigma-Aldrich, #4906845001 and #11836170001, respectively). Cytokine expression was measured with the MSD Mouse Proinflammatory V-Plex Plus Kit (Mesoscale Discovery) and a SECTOR Imager 6000 Plate Reader according to the manufacturer’s instructions. Data were analyzed using the MSD Discovery Workbench software.

### 2.7. Western blot

Hippocampi were homogenized with Tris Lysis buffer (20 mM Tris-HCl, 150 mM NaCl, 1 mM EDTA, 1 mM EGTA, 1% Triton-X-100, pH 7.5) containing phosphatase and protease inhibitors (Sigma-Aldrich, #4906845001 and #11836170001, respectively). Homogenates were sonicated, incubated for 20 min at 4°C in a rotating shaker, and spun at maximum speed for 15 min. Supernatants containing the extracted proteins were collected and stored at −80°C. Proteins were resolved by SDS-PAGE on 4-20% TGX stain-free pre-cast gels (Bio-Rad, #567104), transferred onto nitrocellulose membranes (Bio-Rad, #162-0112), and blocked in 5% non-fat milk in 0.1 M PBS with 0.05% Tween (PBS-T). Membranes were probed overnight at 4°C with primary antibodies against proteins of interest (Table 1), then with horseradish peroxidase (HRP)-conjugated secondary antibodies for 1 h at RT. After incubation with Pico PLUS chemiluminescent substrate (ThermoFisher, #34580) or Femto Maximum Sensitivity Substrate (ThermoFisher, #34096), membranes were imaged with a BioRad ChemiDoc™ Touch Version 1.2.0.12 and quantified with Image Lab Software 5.2.1 (BioRad). Data were normalized to the corresponding β-actin and expressed as percentage ± SEM of the control condition.

**Table 1.** Antibodies for western blot analysis.

| Antigen | Host | Provider | Catalog # | Dilution |
| --- | --- | --- | --- | --- |
| SNAP-25 | Rabbit | Synaptic Systems | 111-002 | 1:1000 |
| Syntaxin 1 | Mouse | Synaptic Systems | 110-111 | 1:1000 |
| Synaptotagmin 1/2 | Rabbit | Synaptic Systems | 105-003 | 1:1000 |
| PSD-93 | Mouse | Novus Biologicals | S18-30 | 1:1000 |
| PSD-95 | Rabbit | Synaptic Systems | 124-002 | 1:1000 |
| GluR1 (GluA1) | Rabbit | Millipore/Upstate | 04-855 | 1:1000 |
| GluR2/3 (GluA2/3) | Rabbit | Millipore/Upstate | 07-598 | 1:1000 |
| NMDAR1 (GluN1) | Rabbit | Abcam | AB109182 | 1:1000 |
| NR2A (GluN2A) | Rabbit | Millipore/Upstate | 07-632 | 1:1000 |
| NR2B (GluN2B) | Rabbit | Millipore/Upstate | 06-600 | 1:1000 |
| GLAST (EAAT1) | Rabbit | Alpha Diagnostic | GLAST11-A | 1:1000 |
| TNFR2 | Rabbit | GeneTex | GTX63066 | 1:500 |
| β-actin | Mouse | Sigma-Aldrich | A5441 | 1:10000 |

### 2.8. Golgi-Cox staining and dendritic spine analysis

Following transcardial perfusion with 0.1 M PBS, brains were dissected out, rinsed in distilled water, cut in half, and stained with the FD Rapid Golgi Stain Kit according to the manufacturer’s protocol (FD NeuroTechnologies). Brains were protected from light and immersed into a 1∶1 mixture of solutions A and B for 14 days at RT, then transferred to Solution C for 3 days at RT. Brains were then frozen on dry ice, cryostat-cut in 120 µm-thick serial sections, and collected onto gelatin-coated slides. Sections were left to air dry overnight at RT while protected from light, then stained according to the manufacturer’s protocol. Briefly, slides were rinsed in distilled water, immersed in solutions D and E for 10 min, rinsed in distilled water, dehydrated in graded ethanol solutions (50%, 75%, 95% and 100%), and cleared in xylene. Slides were finally coverslipped with Permount™ mounting media (ThermoFisher, #SP14-100) and imaged on a Keyence BZ-X800 fluorescence microscope at 100x magnification. In the hippocampus CA1 region, for each animal three cells and three dendritic segments per cell (> 10 µm in length) were imaged. Images were analyzed with Reconstruct software (Harris Lab, Center for Learning and Memory, University of Texas, Austin, TX) as previously described [44] to measure dendritic spine density and classify spine morphology. Analyses were performed by an investigator blinded to the genotype.

### 2.9. Electrophysiological recording of long-term potentiation (LTP)

Following deep anesthesia (3% isoflurane, 70% N_2_O, 30% O_2_, 5 min), adult mice were euthanized, their brains quickly removed, and the hippocampi dissected out. Transverse hippocampal slices (400 μm thick) were prepared using a Vibratome 3000 (Leica) as previously described [45]. Slices were maintained in artificial CSF (aCSF) (125 mM NaCl, 2.5 mM KCl, 1.25 mM NaH_2_PO_4_, 25 mM NaHCO_3_, 10 mM D-glucose, 2 mM CaCl_2_, 1 mM MgCl_2_ saturated with 95% O_2_ and 5% CO_2_) for 1 hr at RT, then moved to a recording chamber where they were continuously perfused in aCSF at a rate of 2.5-3 ml/min at 31°C. Field excitatory postsynaptic potentials (fEPSPs) were recorded in the CA1 stratum radiatum in response to stimulation of the Schaffer collateral pathway as previously described [45]. After the initial slope of the fEPSP was measured, and the input-output (I-O) curve obtained, the stimulation intensity was set at 30-40% of the maximum fEPSP to acquire a baseline recording. Baseline responses were recorded at 0.033 Hz frequency and established for at least 20 min before tetanization. LTP was induced with 4 trains of stimulation (each at 100 Hz for 1 sec, separated by 5 min) at the same intensity used for baseline recordings. Paired-pulse facilitation (PPF) was assessed at the same intensity used for baseline recordings with 12.5-250 msec inter-stimulus intervals.

### 2.10. Primary astrocyte cultures and calcium imaging

Brains from postnatal day 5 WT and *Tnfrsf1b*^-/-^ mouse pups were dissected out and dissociated into single cell suspensions with the Papain Neural Tissue Dissociation Kit (Miltenyi Biotec, #130-092-628). Astrocytes were isolated by MACS separation technology using LS columns (Miltenyi Biotec, #130-042-401) after incubation with astrocyte cell surface antigen-2 (ACSA-2) conjugated magnetic microbeads (Miltenyi Biotec, #130-097-679). ACSA-2^+^ cells were seeded onto poly-D-lysine (Millipore Sigma, #P7280) coated 75 cm^2^ flasks (1×10^6^ cells per flask) and maintained in astrocyte complete medium consisting of DMEM + GlutaMAX (ThermoFisher #10566016) supplemented with 10% heat-inactivated fetal bovine serum (FBS) (Gemini Bio, #900-108), 1% penicillin/streptomycin (ThermoFisher, #15070063), and 0.5% Amphotericin B (ThermoFisher #15290018). Cells were cultured for 4 days, with half volume medium replacement every other day. Cells were then detached with 0.05% Trypsin-EDTA (ThermoFisher, #25300062), and re-plated onto poly-D-lysine coated 15 mm glass coverslips inserted into the wells of a 24-well plate (80,000 cells/well). After 3 days in culture, cells were loaded with the calcium indicator Fluo-4/AM (2.5 um, ThermoFisher Cat# F14201) for 60 min at 37°C. Coverslips were then transferred to a recording chamber and mounted onto an Olympus IX70 microscope equipped with a 10x objective and PCO SensiCam camera (PCO, Pioneer in Cameras and Optoelectronics). Cells were gravity perfused with recording buffer (110 mM NaCl, 5.4 mM KCl, 1.8 mM CaCl_2_, 0.8 mM MgCl_2_, 10 mM D-glucose and 10 mM HEPES at pH 7.4) at a speed of 10 ml/min for 30 sec, then stimulated with ATP (100 μM or 1 mM concentration diluted in recording buffer, Millipore Sigma Cat# A6419-1G). Calcium transients were imaged for 8.5 min and acquired with MicroManager 2.0 software at a frequency of 1 Hz, with a 100 msec exposure time and a spatial resolution of 1024×1024 pixels.

### 2.11. Next generation RNA sequencing

Bulk RNA sequencing was performed on FACS-sorted hippocampal astrocytes collected from adult male *Gfap^creERT2^:Tnfrsf1b^fl/fl^* and *Tnfrsf1b^fl/fl^* mice. Cells were sorted with a FACSAria instrument (BD Biosciences) based on ACSA-2^+^ labeling, collected in lysis buffer, and processed for RNA extraction using the SMART-Seq V4 UltraTM Low Input RNA Kit (Clontech Laboratories), according to the manufacture’s protocol. Library preparation and RNA sequencing were done at the John P. Hussman Institute for Human Genomics (Miller School of Medicine, University of Miami) using an Illumina HiSeq 2500 ultra-high throughput sequencing system. Paired-end, 125 base pairs were generated and analyzed as previously published [40, 46]. Differential gene expression analysis was performed using DESeq2, with normalized gene expression levels represented as fragments per kilobase per million mapped reads (FPKM). For the analysis of differential gene expression, genes with a padj ≤ 0.05 and an absolute log2 fold change (|LogFc|) ≥ 1.5 were included. For Ingenuity Pathway Analysis (IPA), genes with a padj ≤ 0.1 and an absolute log2 fold change (|LogFc|) ≥ 1.0 were included. The accession number for the sequencing data reported in this paper is GEO:GSE288931.

### 2.12. RNA isolation and real-time RT PCR

RNA was extracted from FACS-sorted astrocytes collected from adult male *Gfap^creERT2^:Tnfrsf1b^fl/fl^* and *Tnfrsf1b^fl/fl^* mice using the Arcturus PicoPure RNA isolation kit (Applied Biosystems, # KIT0204) as previously described [47]. Complementary DNA equal to 5 ng of initial total RNA was used as a template in each PCR reaction. Reverse transcription was performed using the SensiScript kit (Qiagen, #205213) according to manufacturer’s protocol. Real-time PCR was run in the QuantStudio 3 Real Time PCR system (Applied Biosystems) with PowerUP SYBR® Green PCR MasterMix (Applied Biosystems). Relative gene expression was calculated with the comparative Ct (ΔΔCt) method after normalization to glyceraldehyde-3-phosphate dehydrogenase (GAPDH) gene expression. Primer sequences for *Tnfrsf1a* were: forward = 5′ gcccgaagtctactccatcatttg 3′; reverse = 5′ ggctggggagggggctggagttag 3′. Primer sequences for *Gapdh* were: forward = 5′ gaggccggtgctgagtatgtcgtg 3′; reverse = 5′ tcggcagaaggggcggagatga 3′.

### 2.13. Statistical Analysis

Statistical analyses were performed with GraphPad Prism software. Details of sample size for each experiment are included in the figure legends. All data were tested for normality with the D’Agostino & Pearson test. Data not normally distributed were analyzed with non-parametric tests. Comparisons of multiple groups in the Sholl analyses were analyzed by two-way ANOVA followed by Sidak test. Single comparisons of normally distributed data were analyzed with Student’s *t* test. Data are expressed as mean ± SEM, and p values equal or less than 0.05 were considered statistically significant.

## 3. Results

### 3.1. *Gfap^creERT2^*:*Tnfrsf1b^fl/fl^* mice are an efficient model to study TNFR2 signaling in astrocytes

To investigate the role of TNFR2 signaling in astrocytes in physiological conditions, we generated *Gfap^creERT2^:Tnfrsf1b^fl/fl^* mice with inducible conditional ablation of the *Tnfrsf1b* gene in astrocytes. *Tnfrsf1b^fl/fl^* mice were used as controls in all experiments. Recombination efficiency and specificity in astrocytes was quantified by crossing *Gfap^creERT2^:Tnfrsf1b^fl/fl^* mice with *Rosa26-Eyfp* reporter mice to obtain *Gfap^creERT2^:Eyfp:Tnfrsf1b^fl/fl^* triple transgenics where, upon tamoxifen administration, *Tnfrsf1b* was ablated and *Eyfp* expressed in astrocytes simultaneously. Recombination efficiency, measured as percentage of GFAP^+^ astrocytes expressing the EYFP protein was robust and comparable in *Gfap^creERT2^:Eyfp:Tnfrsf1b^fl/fl^* and *Gfap^creERT2^:Eyfp* controls, with approximately 75-80% of GFAP^+^ cells expressing EYFP in the hippocampus (Fig. 1A). Furthermore, EYFP was localized only to GFAP^+^ astrocytes, demonstrating specificity (Fig. 1B). Quantification of TNFR2 protein expression in the hippocampus of both male and female mice showed a reduction by about 50% in *Gfap^creERT2^:Tnfrsf1b^fl/fl^* mice compared to *Tnfrsf1b^fl/fl^* controls in the hippocampus (Fig. 1C, K).

**Figure 1.**
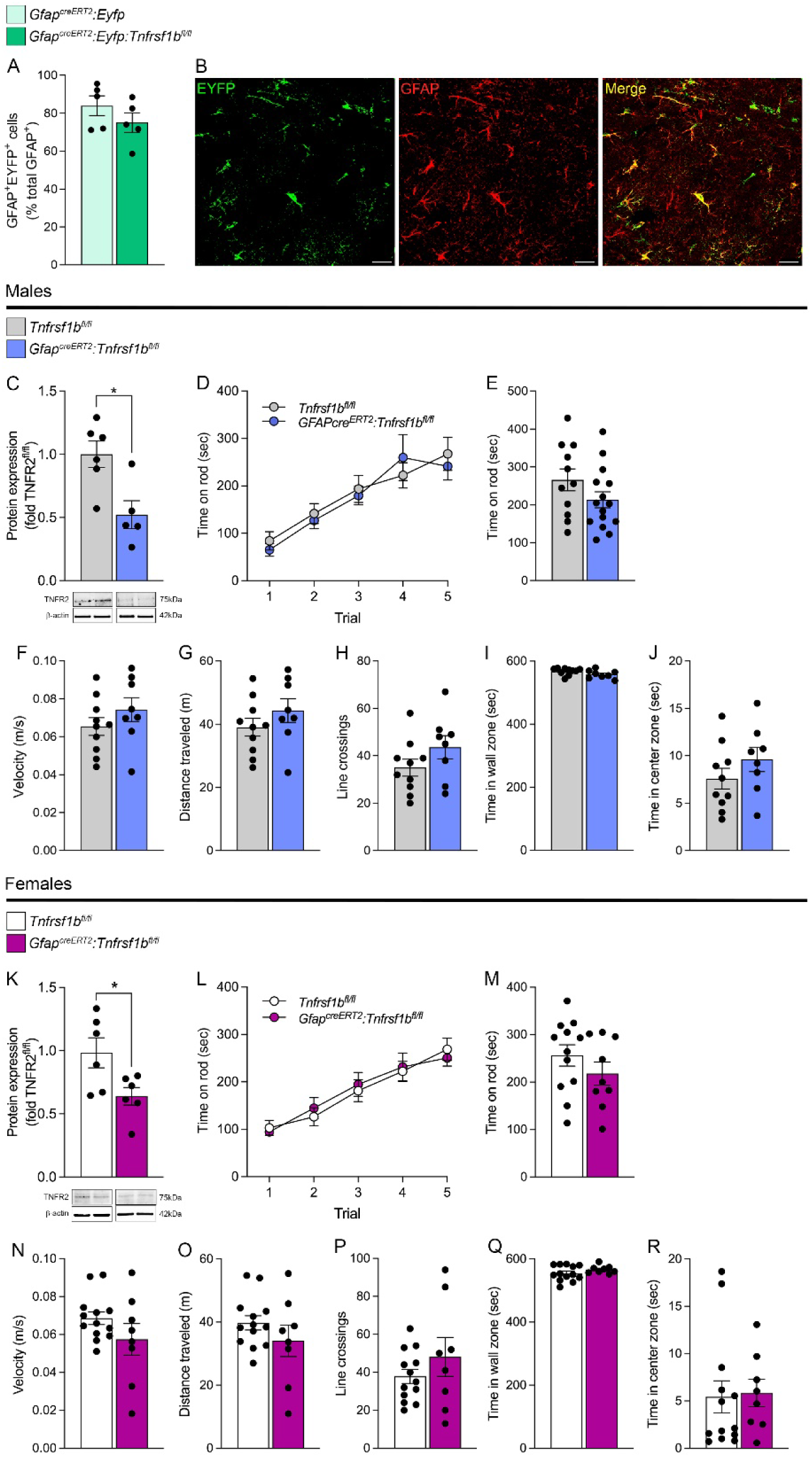
Characterization of *Gfap^creERT2^:Tnfrsf1b^fl/fl^* mice with conditional ablation of the *Tnfrsf1b* gene in astrocytes. (A-B) Cre recombination efficiency assessed in *Gfap^creERT2^:Eyfp:Tnfrsf1b^fl/fl^* mice with ablation of the *Tnfrsf1b* gene and concomitant *Eyfp* reporter expression in comparison with *Gfap^creERT2^:Eyfp* reporter mice. (A) Stereological quantification of recombined GFAP^+^EYFP^+^ cells in the hippocampus expressed as % of total GFAP^+^ cells. (B) Representative images showing colocalization of EYFP immunolabeling (green) with the astrocyte-specific marker GFAP (red) in the hippocampus of *Gfap^creERT2^:Eyfp:Tnfrsf1b^fl/fl^* mice; scale bar = 50 μm. (C, K) Western blot analysis of TNFR2 in the hippocampus of *Tnfrsf1b^fl/fl^* and *Gfap^creERT2^:Tnfrsf1b^fl/fl^* male (C) and female (K) mice; n=5-6/group, *p≤0.05, Student’s *t* test. Results represent average ± SEM. (D, E, L, M) Assessment of locomotor coordination with the rotarod test in male (D, E) and female (L, M) *Tnfrsf1b^fl/fl^* and *Gfap^creERT2^:Tnfrsf1b^fl/fl^* mice: time spent on the rod during a 5-day pre-training session (D, L) and during the test day (E, M). (F-J, N-R) Assessment of locomotor activity with the open field test in *Tnfrsf1b^fl/fl^* and *Gfap^creERT2^:Tnfrsf1b^fl/fl^* male and female mice: measures of velocity (F, N), total distance traveled (G, O), number of line crossings (H, P), time spent in wall zone (I, Q),and time spent in center zone (J, R) were obtained; n=8-15/group. Results represent average ± SEM.

To determine if ablation of *Tnfrsf1b* in astrocytes interfered with basal locomotor function, *Gfap^creERT2^:Tnfrsf1b^fl/fl^* mice were assessed with rotarod (Fig. 1D-E, L-M) and open field (Fig. 1F-J, N-R) tests in comparison to *Tnfrsf1b^fl/fl^* controls. Neither male nor female mice exhibited locomotor dysfunction. Indeed, coordination, evaluated as the time mice remained on the rotating rod, was comparable between *Gfap^creERT2^:Tnfrsf1b^fl/fl^* and *Tnfrsf1b^fl/fl^* mice (Fig. 1D, E, L, M). Likewise, locomotion and exploratory behavior were not affected as *Tnfrsf1b* ablated mice traveled similar distances and with the same velocity as controls in the open field (Fig. 1F, G, N, O). Additionally, we analyzed anxiety-like behaviors with the light-dark transition test and uncovered significant differences between *Gfap^creERT2^:Tnfrsf1b^fl/fl^* and *Tnfrsf1b^fl/fl^* mice. Both males and females exhibited a reduction in distance traveled, total movement time, time spent in the light zone, and frequency of wall explorations, which was paralleled by an increase in latency to light zone exploration (Fig. 2). This suggests that astroglial TNFR2 may play a role in regulating the neural networks implicated in stress- and anxiety-related behaviors, which are controlled not only by the hippocampus but also by the cortex and amygdala.

**Figure 2.**
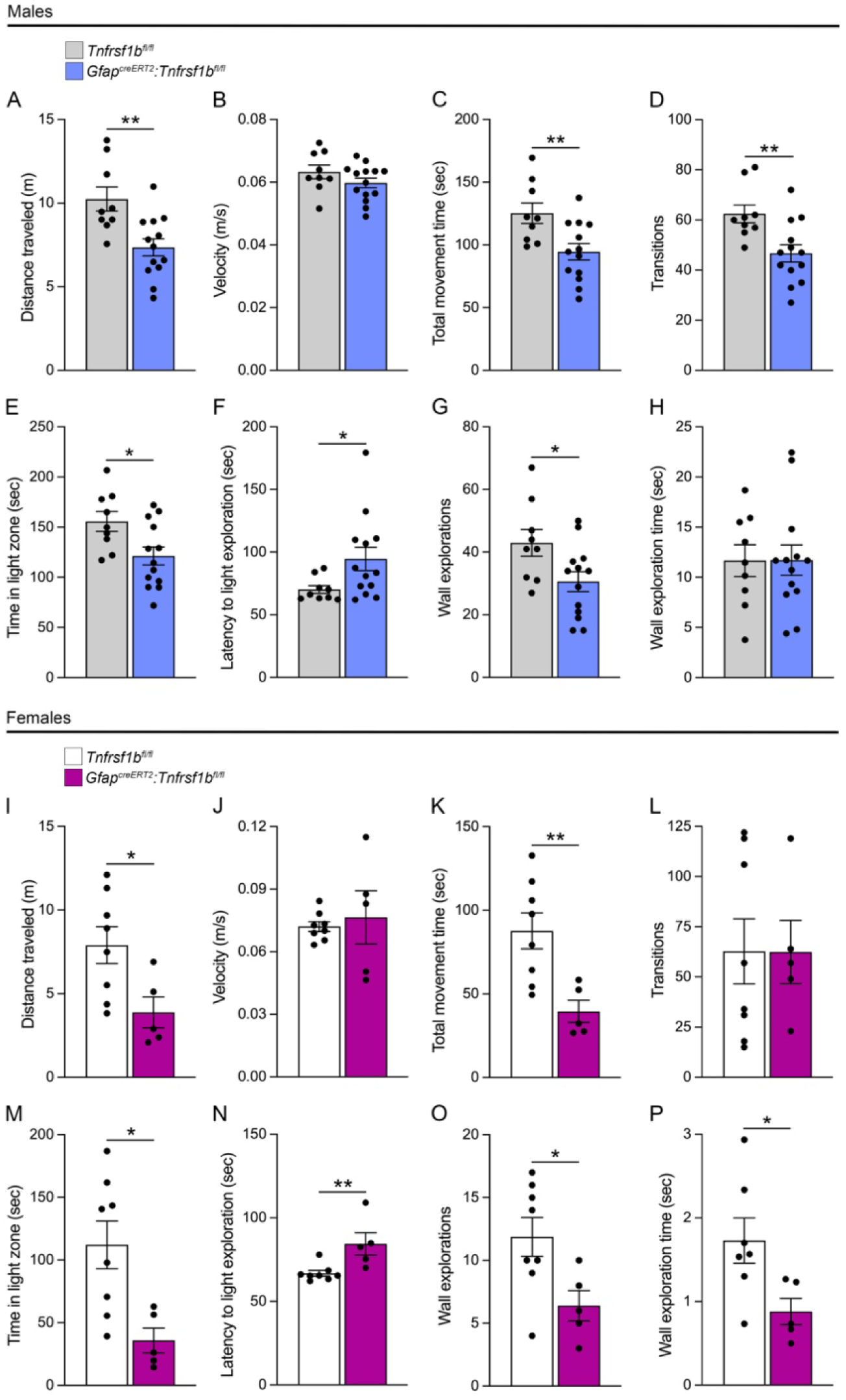
Ablation of astroglial TNFR2 results in anxiety-type behavior. Assessment of anxiety-based behavior with the light-dark transition test in *Tnfrsf1b^fl/fl^* and *Gfap^creERT2^:Tnfrsf1b^fl/fl^* male (A-H) and female (I-P) mice: (A, I) total distance traveled, (B, J) velocity, (C, K) total movement time, (D, L) number of transitions between chambers, (E, M) time spent in the light chamber, (F, N) latency to light chamber exploration, (G, O) number of wall explorations, and (H, P) wall exploration time were measured; n=5-13/group, *p≤0.05, **p≤0.01, Student’s *t* test. Results represent average ± SEM.

### 3.2. TNFR2 ablation in astrocytes alters the expression of hippocampal proteins involved in synaptic transmission and plasticity

To unravel the role of astroglial TNFR2 in the regulation of synaptic function, in the hippocampus of *Gfap^creERT2^:Tnfrsf1b^fl/fl^* and *Tnfrsf1b^fl/fl^* male and female mice we quantified the expression of proteins implicated in synaptic transmission and plasticity, specifically pre- and post-synaptic molecules, glutamate receptors, and glutamate transporters (Fig. 3).

**Figure 3.**
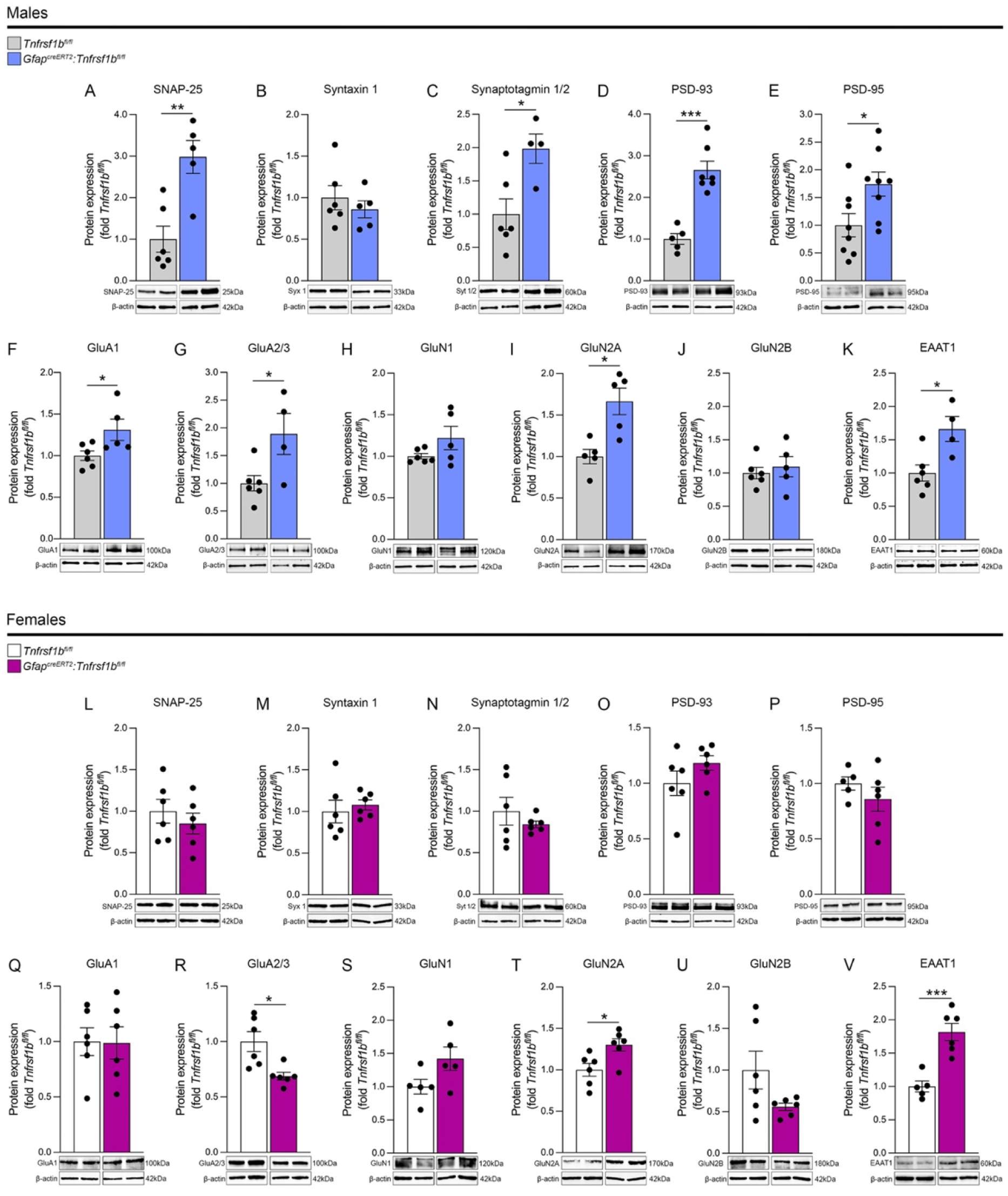
Ablation of astroglial TNFR2 alters the expression of hippocampal synaptic proteins, glutamate receptors and transporters. Western blot analysis of SNAP-25 (A, L), Syntaxin 1 (B, M), Synaptotagmin 1/2 (C, N), PSD-93 (D, O), PSD-95 (E, P), GluA1 (F, Q), GluA2/3 (G, R), GluN1 (H, S), GluN2A (I, T), GluN2B (J, U), and EAAT1 (K, V) in the hippocampus of male (A-K) and female (L-V) *Tnfrsf1b^fl/fl^* and *Gfap^creERT2^:Tnfrsf1b^fl/fl^* mice; n=4-8/group, *p≤0.05, **p≤0.01, ***p≤0.001, Student’s *t* test. Results are expressed as fold of corresponding controls and represent average ± SEM.

Among pre-synaptic SNARE complex proteins that are essential for vesicle fusion and neurotransmitter release [48], we observed a robust upregulation of SNAP-25 and Synaptotagmin1/2 in male *Gfap^creERT2^:Tnfrsf1b^fl/fl^* mice compared to *Tnfrsf1b^fl/fl^* controls (Fig. 3A, C), but no change in Syntaxin 1 (Fig. 3B). This was paralleled by the upregulation of the post-synaptic scaffold proteins PSD-93 and PSD-95 (Fig. 3D, E). Notably, none of these pre- and post-synaptic proteins was altered in female *Gfap^creERT2^:Tnfrsf1b^fl/fl^* mice compared to *Tnfrsf1b^fl/fl^* control mice (Fig. 3L-N, O, P).

Next, we quantified the expression of NMDA and AMPA glutamate receptor subunits, which are both crucial for synaptic plasticity. AMPA receptor subunits GluA1 and GluA2/3 were upregulated in male *Gfap^creERT2^:Tnfrsf1b^fl/fl^* mice compared to *Tnfrsf1b^fl/fl^* controls (Fig. 3F, G). This did not occur in female *Gfap^creERT2^:Tnfrsf1b^fl/fl^* mice, where GluA1 remained the same (Fig. 3Q) and GluA2/3 was downregulated (Fig. 3R) compared to *Tnfrsf1b^fl/fl^* controls. The expression pattern of NMDA receptor subunits was similar between male and female mice, with no changes in GluN1 and GluN2B (Fig. 3H, J, S, U), but with a significant upregulation of GluN2A (Fig. 3I, T) in *Gfap^creERT2^:Tnfrsf1b^fl/fl^* mice compared to *Tnfrsf1b^fl/fl^* controls.

Finally, we quantified the expression of the astrocytic glutamate transporter EAAT1, which is responsible for glutamate reuptake from the synaptic cleft, and found it to be upregulated in both male and female *Gfap^creERT2^:Tnfrsf1b^fl/fl^* mice compared to *Tnfrsf1b^fl/fl^* controls (Fig. 3K, V).

Collectively, these data point at a key role for astroglial TNFR2 in physiologic synaptic transmission and plasticity in the hippocampus through the regulation of pre-and post-synaptic mechanisms. Such function is sex-dependent, with a more pronounced role for TNFR2 in males versus females.

### 3.3. TNFR2 ablation in astrocytes does not alter neuronal arborization and dendritic spine morphology in the hippocampus

The observed changes in neuronal proteins implicated in synaptic transmission and plasticity prompted us to test whether ablation of astroglial TNFR2 might also affect neuronal structure and numbers in the hippocampus. When we quantified the number of NeuN^+^ neurons in male mice, we found no differences in the CA1 and CA3 regions, but a mild increase in the dentate gyrus (DG) of *Gfap^creERT2^:Tnfrsf1b^fl/fl^* mice compared to *Tnfrsf1b^fl/fl^* controls (Fig. 4A), suggesting a potential effect on hippocampal neurogenesis. In contrast, we found no changes in neuronal numbers in female *Gfap^creERT2^:Tnfrsf1b^fl/fl^* mice compared to controls (Fig. 4G). Next, we evaluated axonal and dendritic arborization complexity by Sholl analysis and observed no differences between *Gfap^creERT2^:Tnfrsf1b^fl/fl^* and *Tnfrsf1b^fl/fl^* mice, both males and females. Indeed, number of process intersections at increasing distance from the nucleus (measure of ramification), number of processes, total process length, and radius at which the maximum number of intersections was measured (Radius w/ max intersections) were all comparable between genotypes (Fig. 4B-E, H-K). Lastly, we evaluated dendritic spine morphology following Golgi straining by quantifying the percentage of mushroom shaped and thin/long-thin spines which have been associated with mature/strong and damaged/weak synaptic connections, respectively [44]. We found no differences in spine morphology between *Gfap^creERT2^:Tnfrsf1b^fl/fl^* and *Tnfrsf1b^fl/fl^* mice for both sexes (Fig. 4F, L).

**Figure 4.**
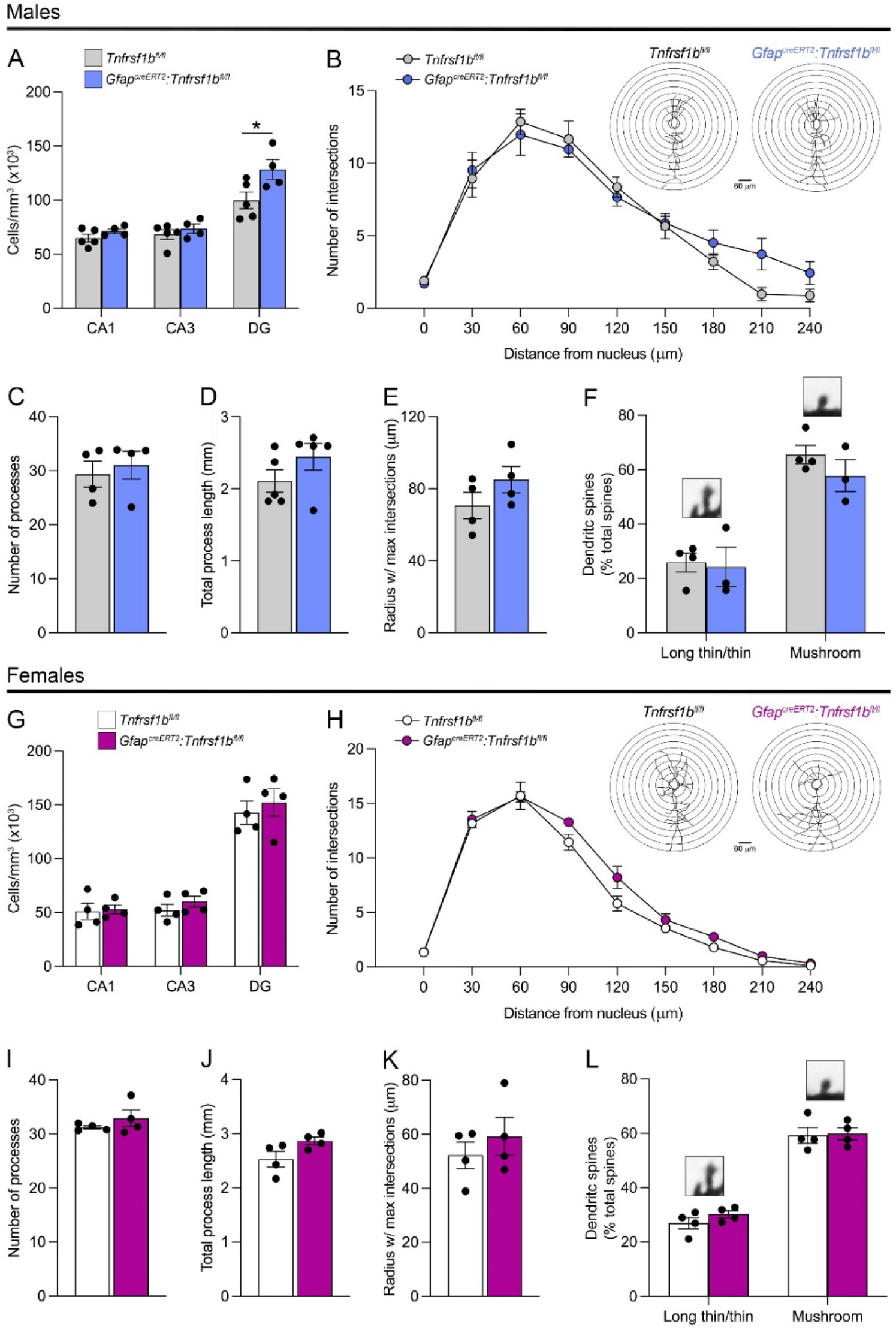
Ablation of astroglial TNFR2 does not alter the morphology of hippocampal neurons. (A, G) Stereological quantification of NeuN^+^ neurons in CA1, CA3, and DG of male (A) and female (G) *Tnfrsf1b^fl/fl^* and *Gfap^creERT2^:Tnfrsf1b^fl/fl^* mice. (B-E, H-K) Sholl analysis of neuronal arborization in male (B-E) and female (H-K) *Tnfrsf1b^fl/fl^* and *Gfap^creERT2^:Tnfrsf1b^fl/fl^* mice: (B, H) number of process intersections at increasing distance from the cell nucleus (with representative Sholl traces), (C, I) total process number, (D, J) total process length, and (E, K) radius at which the maximum number of intersections is observed in NeuN^+^ CA1 neurons. (F, L) Analysis of dendritic spine morphology after Golgi staining: quantification of long thin/thin and mushroom-shaped spines in male (F) and female (L) *Tnfrsf1b^fl/fl^* and *Gfap^creERT2^:Tnfrsf1b^fl/fl^* mice; n=4-5/group, *p≤0.05 Student’s *t* test. Results represent average ± SEM.

Together, our data indicate that astroglial TNFR2 does not play a role in neuronal survival, nor impacts neuronal morphology, both in terms of process arborization and spine phenotype. Yet, it may influence DG neurogenesis in males, further pointing at a more pronounced role in male than female mice.

### 3.4. TNFR2 ablation in astrocytes leads to hippocampal astrogliosis in male mice

To examine the impact of astroglial TNFR2 ablation on hippocampal astrocytes, we quantified the number of GFAP^+^ cells in the CA1, CA3, and DG. In male *Gfap^creERT2^:Tnfrsf1b^fl/fl^* mice we found GFAP^+^ astrocytes to be significantly increased in all regions compared to *Tnfrsf1b^fl/fl^* controls (Fig. 5A, B). Sholl analysis in the CA1 uncovered that astrocytes had a higher morphological complexity than control cells (Fig. 5A, bottom panels), demonstrated by a significant increase in the number of process intersections at a distance of 8 μm from the cell nucleus (Fig. 5C), and an overall trend of increased intersection numbers at all distances. Furthermore, compared to *Tnfrsf1b^fl/fl^* controls, astrocytes in male *Gfap^creERT2^:Tnfrsf1b^fl/fl^* mice had a higher number of cell processes and a higher radius at which the maximum number of intersections was observed (Radius w/ max intersections, Fig. 5D), although no differences were found in the average and total process length (Fig. 5D). Notably, none of these changes were recapitulated in females where *Gfap^creERT2^:Tnfrsf1b^fl/fl^* mice had the same number of astrocytes as *Tnfrsf1b^fl/fl^* controls (Fig. 5E, F) as well as similar cell complexity, including process number and length (Fig. 5E, G, H).

**Figure 5.**
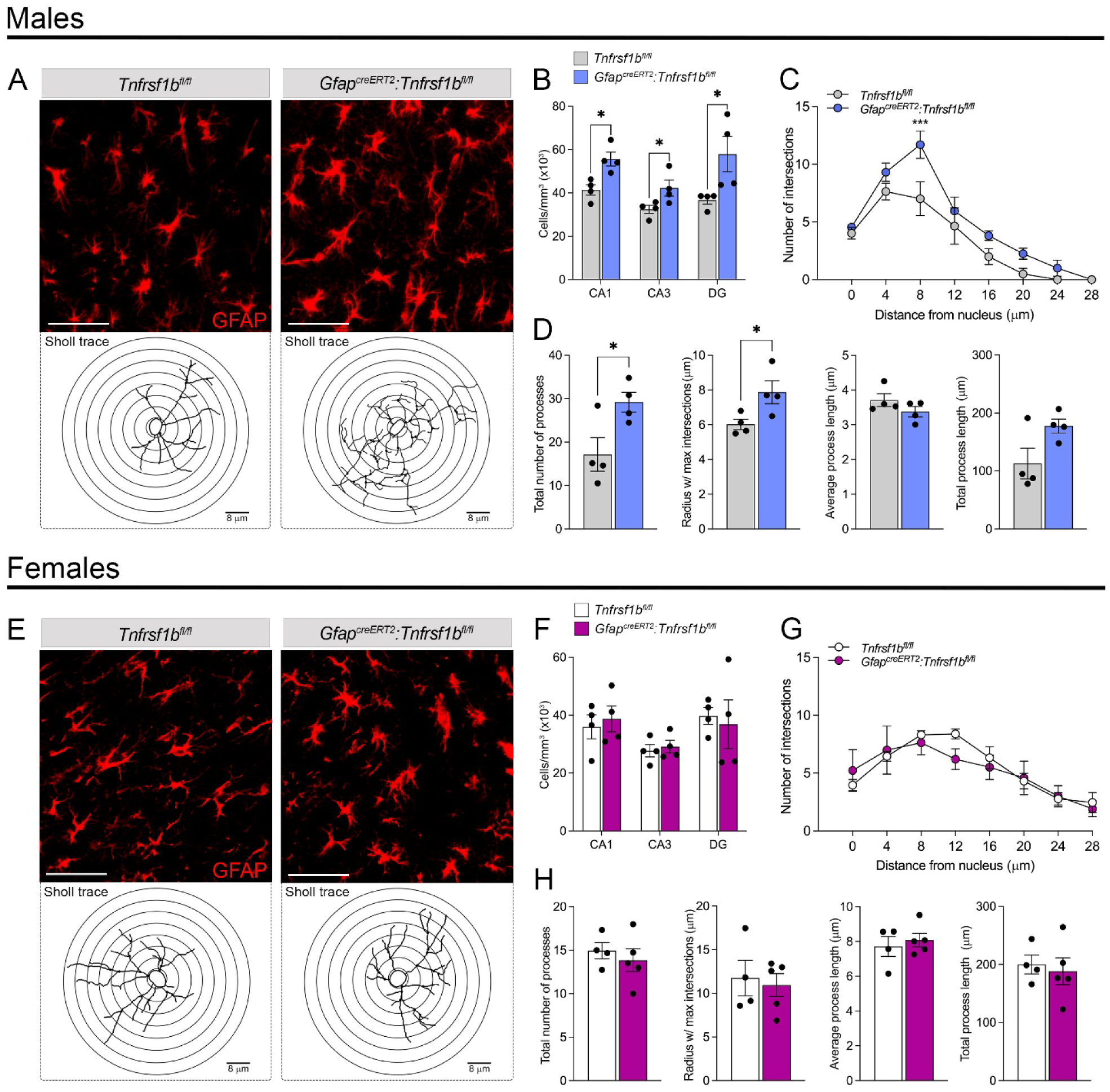
Ablation of astroglial TNFR2 leads to hippocampal astrogliosis in male mice. (A, E) Representative images of GFAP^+^ astrocytes with corresponding Sholl traces in the CA1 region of male (A) and female (E) *Tnfrsf1b^fl/fl^* and *Gfap^creERT2^:Tnfrsf1b^fl/fl^* mice; scale bar = 50 μm. (B, F) Stereological quantification of GFAP^+^ astrocytes in CA1, CA3, and DG of male (B) and female (F) mice. (C, D, G, H) Sholl analysis of astrocyte morphology in the CA1 region: (C, G) number of process intersections at increasing distance (4 μm increments) from the nucleus, (D, H) total number of processes, radius at which the maximum number of intersections is observed, average process length, and total process length; n=4-5/group, *p≤0.05 Student’s *t* test, ***p≤0.001, two-way ANOVA, Sidak’s test. Results represent average ± SEM.

Collectively, these data point at a male-specific astrogliotic response in the absence of TNFR2, suggesting that TNFR2 plays a role in controlling the proliferation and activation state of astrocytes in physiological conditions in a sex-specific manner.

### 3.5. TNFR2 ablation in astrocytes leads to hippocampal microgliosis in male mice

To examine the impact of astroglial TNFR2 ablation on hippocampal microglia, we quantified the number of Iba1^+^ cells in the CA1, CA3, and DG. In male *Gfap^creERT2^:Tnfrsf1b^fl/fl^* mice we found Iba^+^ microglia to be significantly increased (almost doubled) in all regions compared to *Tnfrsf1b^fl/fl^* controls (Fig. 6A, B). Morphologically, these cells did not exhibit any differences from microglia in control mice (Fig. 6A, bottom panels; Fig. 6C, D), displaying similar process ramifications, length, and numbers. Microglia in female *Gfap^creERT2^:Tnfrsf1b^fl/fl^* mice were comparable to those in *Tnfrsf1b^fl/fl^* controls, both in number and morphology (Fig. 6E-H).

**Figure 6.**
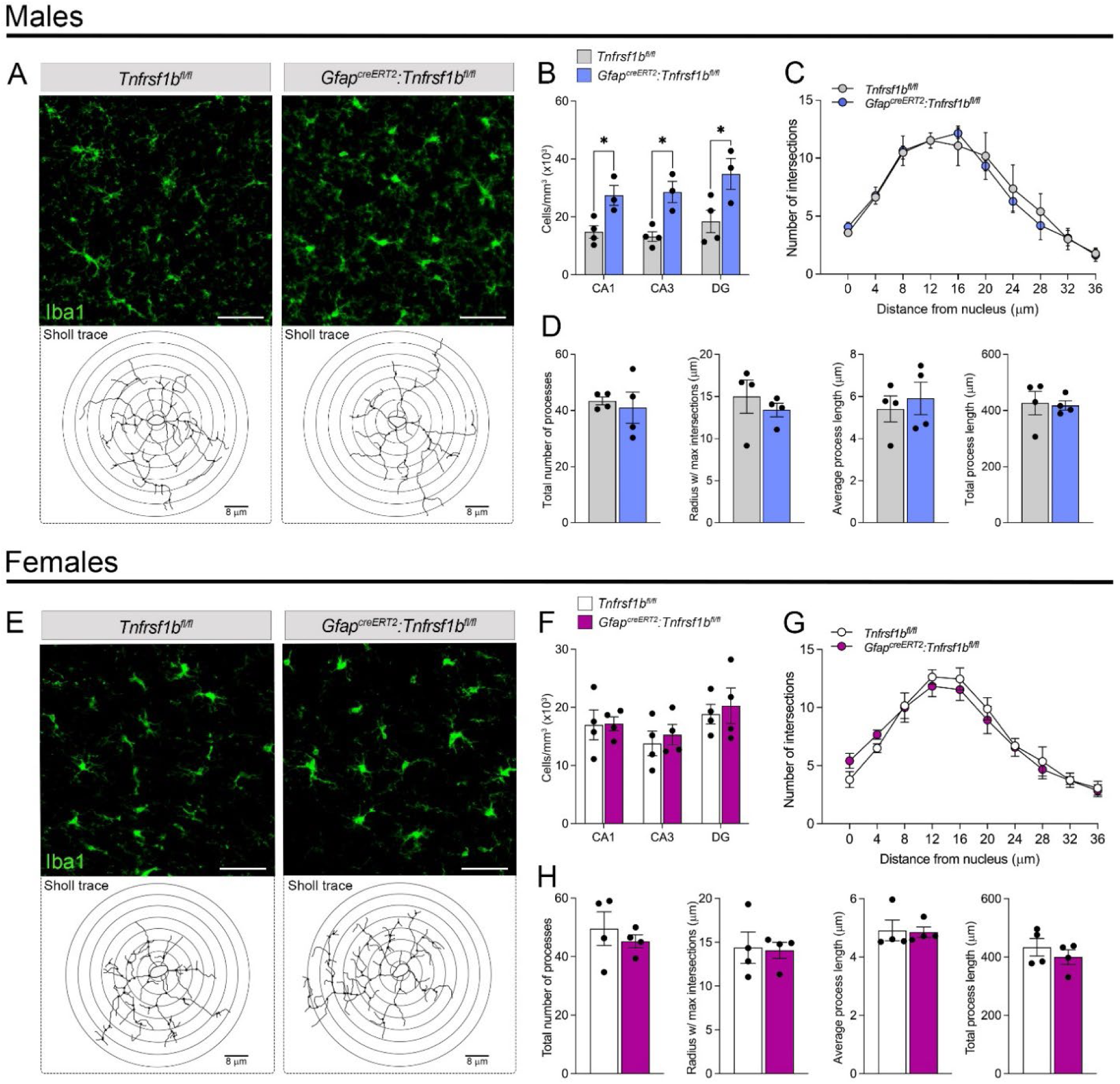
Ablation of astroglial TNFR2 leads to hippocampal microgliosis in male mice. (A, E) Representative images of Iba1^+^ microglia with corresponding Sholl traces in the CA1 region of male (A) and female (E) *Tnfrsf1b^fl/fl^* and *Gfap^creERT2^:Tnfrsf1b^fl/fl^* mice; scale bar = 50 μm. (B, F) Stereological quantification of Iba^+^ microglia in CA1, CA3, and DG of male (B) and female (F) mice. (C, D, G, H) Sholl analysis of microglia morphology in the CA1 region: (C, G) number of process intersections at increasing distance (4 μm increments) from the nucleus, (D, H) total number of processes, radius at which the maximum number of intersections is observed, average process length, and total process length; n=3-4/group, *p≤0.05 Student’s *t* test. Results represent average ± SEM.

Our data point at a male-specific hyperplasic response of microglia in the absence of astroglial TNFR2, suggesting that, similar to astrocytes, TNFR2 is implicated in the control of microglia proliferation in physiological conditions.

### 3.6. TNFR2 ablation in astrocytes results in increased glial reactivity

To further probe for glial reactivity in the hippocampus upon astroglial TNFR2 ablation, we quantified the amount of [^3^H]PBR28, which selectively binds to TSPO, in brain tissue sections of male and female *Gfap^creERT2^:Tnfrsf1b^fl/fl^* and *Tnfrsf1b^fl/fl^* mice (Fig. 7A-E, G-K). Since TSPO is expressed by activated microglia and astrocytes [49] [50], this approach allows for a quantitative measure of glial reactivity. Indeed, TSPO binding with selective radioligands is the only validated PET imaging modality to quantify microglial reactivity *in vivo* [51]. In male mice, we detected a significant increase in TSPO binding in whole brain sections, in the hippocampus, cortex, and thalamus of *Gfap^creERT2^:Tnfrsf1b^fl/fl^* mice compared to *Tnfrsf1b^fl/fl^* controls (Fig. 7B-E), with the highest TSPO signal localized to the hippocampus (Fig. 7A, C). This further reinforces that astroglial TNFR2 ablation in male mice leads to increased glial reactivity in the hippocampus and indicates that other brain regions are also impacted. Conversely, female *Gfap^creERT2^:Tnfrsf1b^fl/fl^* mice did not display differences compared to corresponding *Tnfrsf1b^fl/fl^* controls although, interestingly, female mice had overall more TSPO binding compared to male mice (Fig. 7G-L), further highlighting the sex differences in astroglial TNFR2 signaling.

**Figure 7.**
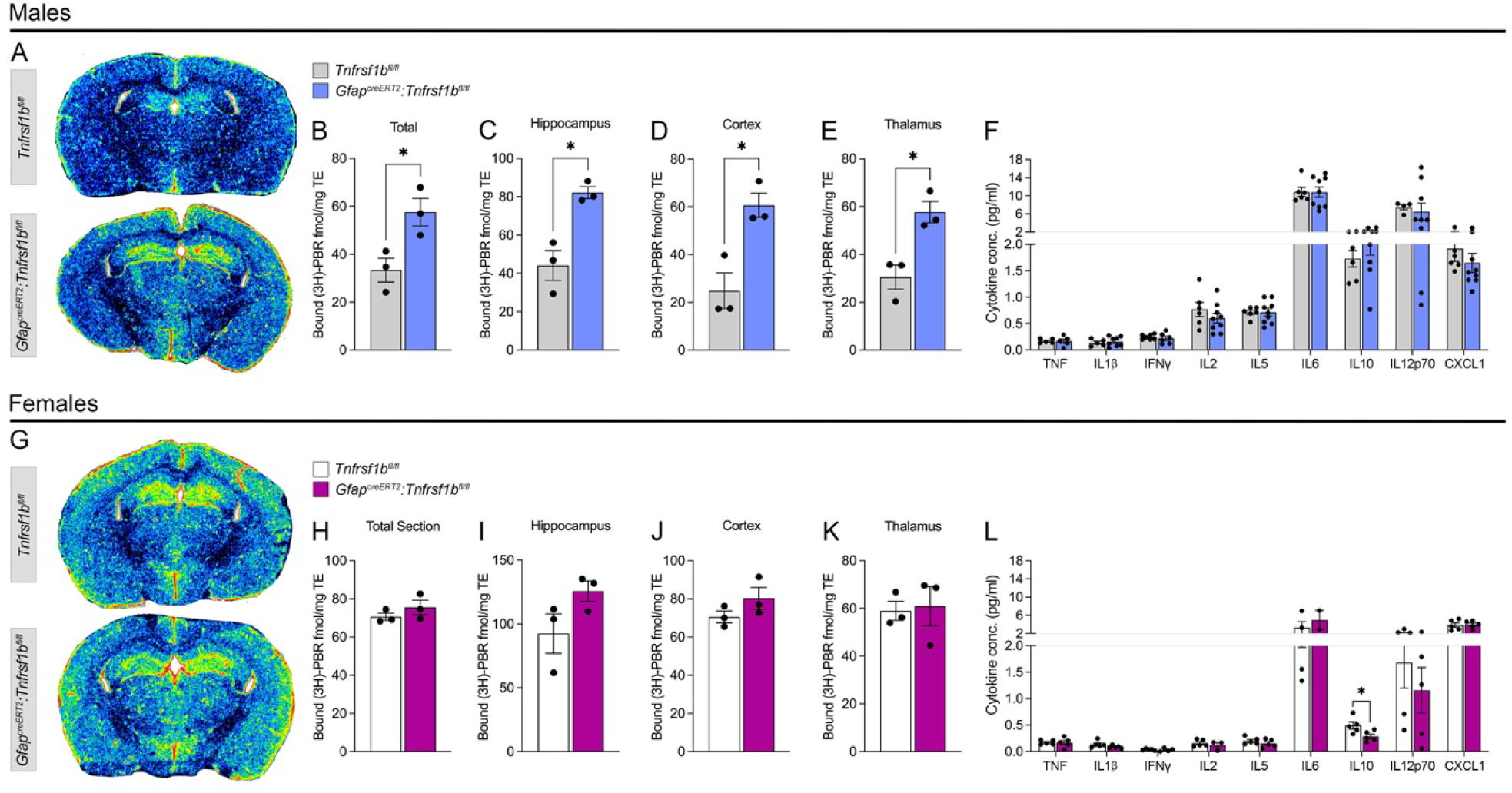
Ablation of astroglial TNFR2 results in increased TSPO binding in the hippocampus, cortex and thalamus of male mice. (A, G) Representative autoradiograms depicting [^3^H]PBR28 binding to TSPO in brain sections from male (A) and female (B) *Tnfrsf1b^fl/fl^* and *Gfap^creERT2^:Tnfrsf1b^fl/fl^* mice. (B-E, H-K) Quantification of [^3^H]PBR28 binding in male (B-E) and female (H-K) *Tnfrsf1b^fl/fl^* and *Gfap^creERT2^:Tnfrsf1b^fl/fl^* mice. Data are expressed as fmol/mg of tissue. n=3/group, *p≤0.05 Student’s *t* test. Results represent average ± SEM. (F, L) Quantification of cytokines and chemokines in the hippocampus of male (F) and female (L) *Tnfrsf1b^fl/fl^* and *Gfap^creERT2^:Tnfrsf1b^fl/fl^* mice by multiplex assay. Data are expressed as pg/mg of tissue, n=5-9/group, *p≤0.05 Student’s *t* test. Results represent average ± SEM.

To determine whether the increased hippocampal glial reactivity in male mice lacking astroglial TNFR2 was associated with changes in inflammatory/immunomodulatory molecules, we assessed cytokine and chemokine protein expression in the hippocampus by multiplex analysis (Fig. 6F, L). Notably, no differences were observed between male *Gfap^creERT2^:Tnfrsf1b^fl/fl^* mice and corresponding *Tnfrsf1b^fl/fl^* controls (Fig. 7F). In female mice, only IL10 was differentially expressed and found to be significantly lower in *Gfap^creERT2^:Tnfrsf1b^fl/fl^* mice compared to *Tnfrsf1b^fl/fl^* controls (Fig. 7L). Yet, this reduction in IL10, an immunosuppressive and anti-inflammatory cytokine with known effects on glial activation [52], did not influence glial reactivity in female mice.

Collectively, these data indicate that astroglial TNFR2 has a role in containing glial reactivity primarily in male mice. Since the shift from homeostatic to reactive glia profoundly affects neuronal circuit functionality [53, 54], our data provide further evidence that astroglial TNFR2 is essential for maintaining homeostatic glia-neuron and glia-glia communication at the basis of synaptic function.

### 3.7. Ablation of TNFR2 in astrocytes impairs hippocampal synaptic plasticity and cognitive function

Since male *Gfap^creERT2^:Tnfrsf1b^fl/fl^* mice exhibited the strongest alterations in hippocampal proteins involved in synaptic function combined with the highest increase in hippocampal glial reactivity, we tested whether these changes translated into functional alterations of synaptic plasticity by evaluating hippocampal LTP. Compared to *Tnfrsf1b^fl/fl^* controls, *Gfap^creERT2^:Tnfrsf1b^fl/fl^* male mice showed suppressed LTP (Fig. 8A), with fEPSP slopes dropping rapidly to pre-stimulation levels after tetanization.

**Figure 8.**
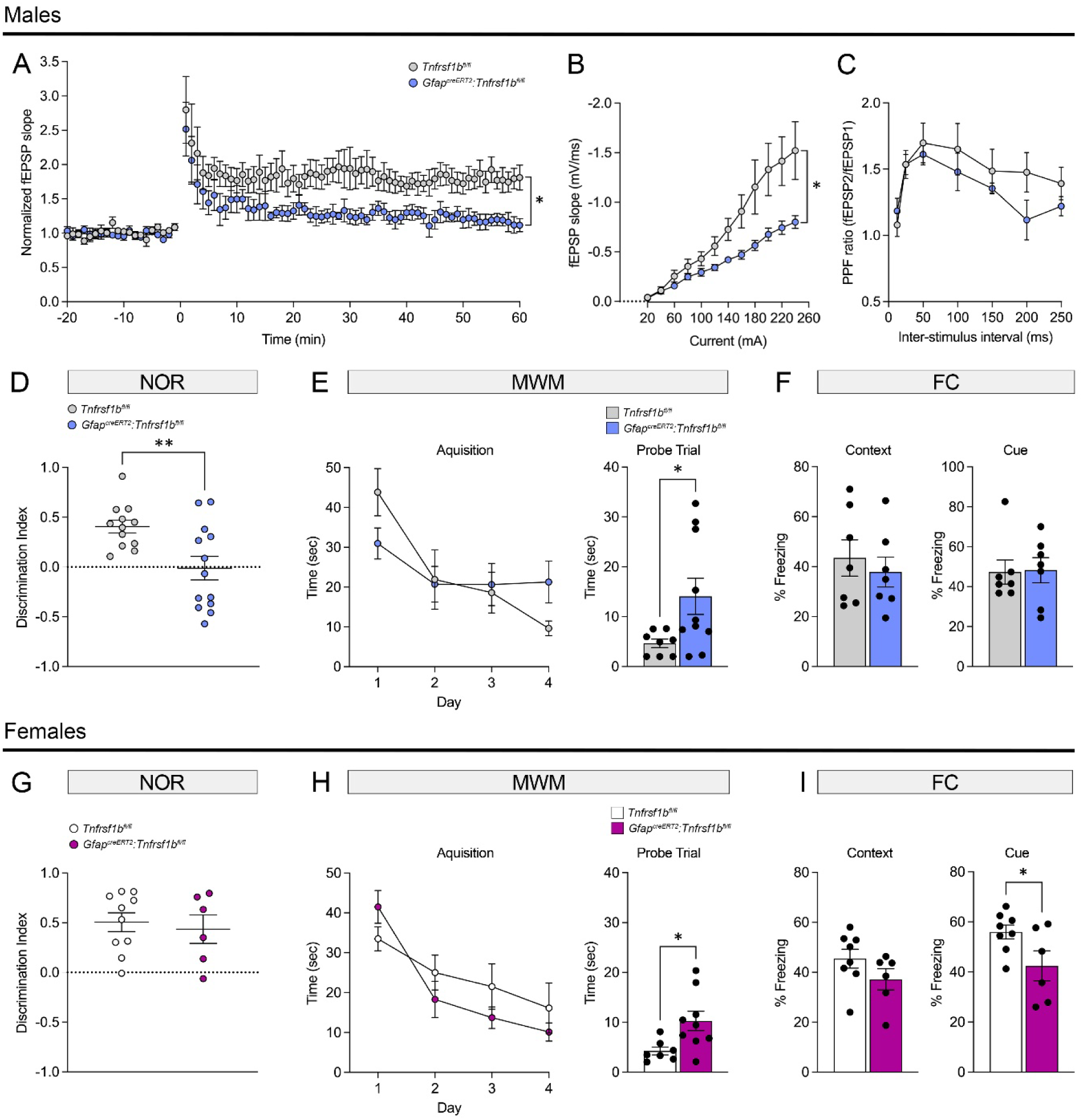
Ablation of astroglial TNFR2 suppresses hippocampal LTP and causes cognitive impairment. (A) Field excitatory postsynaptic potentials (fEPSPs) recorded in the CA1 stratum radiatum after stimulation of the Schaffer collateral pathway in the CA3 of *Tnfrsf1b^fl/fl^* and *Gfap^creERT2^:Tnfrsf1b^fl/fl^* male mice. LTP was induced with 4 trains of 1×100 Hz (1 sec, 5 min intervals) at test stimulation intensity. (B) Input/output (I/O) curves obtained by plotting the field excitatory postsynaptic potential (fEPSP) slopes at increasing stimulus intensities (from 0 to 240 µA in 40 µA steps). (C) Paired-pulse facilitation (PPF) ratio assessed at test stimulation intensity with 12.5-250 msec inter-stimulus intervals; n=7/group, *p≤0.05, repeated measures 2-way ANOVA, Tukey test. Results represent average ± SEM. (D, G) Novel object recognition (NOR) test in male (D) and female (G) *Tnfrsf1b^fl/fl^* and *Gfap^creERT2^:Tnfrsf1b^fl/fl^* mice. The discrimination index (Time with novel object - Time with familiar object / Total time) was recorded as a measure of long-term memory retention. (E, H) Morris water maze (MWM) test in male (E) and female (H) *Tnfrsf1b^fl/fl^* and *Gfap^creERT2^:Tnfrsf1b^fl/fl^* mice. The time to reach the hidden platform during training (acquisition) was quantified as measure of spatial learning, and the time to reach the target area on trial day (probe trial) was quantified as measure of spatial memory. (F, I) Fear conditioning (FC) test in male (F) and female (I) *Tnfrsf1b^fl/fl^* and *Gfap^creERT2^:Tnfrsf1b^fl/fl^* mice. The % freezing response (percent of total time spent freezing) was recorded as measure of context and cue based associative memory; n=6-13/group, *p≤0.05, **p≤0.01, Student’s *t* test. Results represent average ± SEM.

Furthermore, I/O responses shifted downward suggesting depression of basal transmission (Fig. 8B). PPF ratio exhibited a declining trend but was not significantly different between *Gfap^creERT2^:Tnfrsf1b^fl/fl^* and *Tnfrsf1b^fl/fl^* mice (Fig. 8C). Collectively, these data suggest that astroglial TNFR2 is important for maintenance of proper synaptic plasticity in physiological conditions.

Since synaptic plasticity is at the basis of cognitive function, we next evaluated cognitive performance through a series of behavioral tests, specifically novel object recognition (NOR), Morris water maze (MWM) and fear conditioning (FC), both in male and female *Gfap^creERT2^:Tnfrsf1b^fl/fl^* and *Tnfrsf1b^fl/fl^* mice (Fig. 8D-I). Male (Fig. 8D) but not female (Fig. 8G) *Gfap^creERT2^:Tnfrsf1b^fl/fl^* mice had impaired long-term memory, as indicated by their reduced ability to discriminate a novel from a familiar object in the NOR test. Both male and female *Gfap^creERT2^:Tnfrsf1b^fl/fl^* mice showed significant deficits in spatial memory, as demonstrated by the increased time taken to reach the target area during the probe trial of the MWM test (Fig. 8E, H). Finally, while no deficits were observed in context-based associative learning/memory in either sex in the FC test (Fig. 8F, I), female *Gfap^creERT2^:Tnfrsf1b^fl/fl^* mice exhibited deficits in cue-based associative learning/memory (Fig. 8I). Since contextual processing in the FC test has been shown to be mediated by the hippocampus, and cue processing by the amygdala [55], these results suggest that astroglial TNFR2 ablation in female mice may also impair amygdala-dependent cognitive function. Together, these findings demonstrate that astroglial TNFR2 has an important role in cognitive function, both in males and females.

### 3.8. TNFR2 regulates intracellular Ca^2+^ transients in astrocytes

To shed light on the mechanisms by which TNFR2 in astrocytes regulates synaptic function/plasticity and cognition, we evaluated whether TNFR2 played a role in modulating intracellular Ca^2+^ dynamics in astrocytes, given that Ca^2+^ signaling is a key mode of cell-to-cell communication for astrocytes [56], where transient elevations in calcium levels trigger the release of gliotransmitters (e.g. Glutamate, D-serine, ATP, GABA) [57, 58]. These then act on neurons, thereby making astrocytes powerful regulators of neurotransmission and synaptic plasticity. On this basis, we measured intracellular Ca^2+^ ([Ca^2+^]_i_) waves via calcium imaging in *Wt* and *Tnfrsf1b^-/-^* primary mouse astrocytes. Astrocytes lacking TNFR2 (*Tnfrsf1b^-/-^*) exhibited abnormal, spontaneously elevated and intermittent [Ca^2+^]_i_ under basal conditions (Fig. 9A, B, C). Abnormal [Ca^2+^]_i_ peaks were further elevated and prolonged following ATP stimulation (Fig. 9D-K). These findings provide evidence that TNFR2 signaling plays a role in fine-tuning astrocyte-dependent regulatory mechanisms of synaptic function under physiological conditions. As elevation in [Ca^2+^]_i_ is the critical signal for gliotransmitter release, these data suggest that lack of TNFR2 in astrocytes may lead to aberrant release of gliotransmitters, negatively impacting neuronal function.

**Figure 9.**
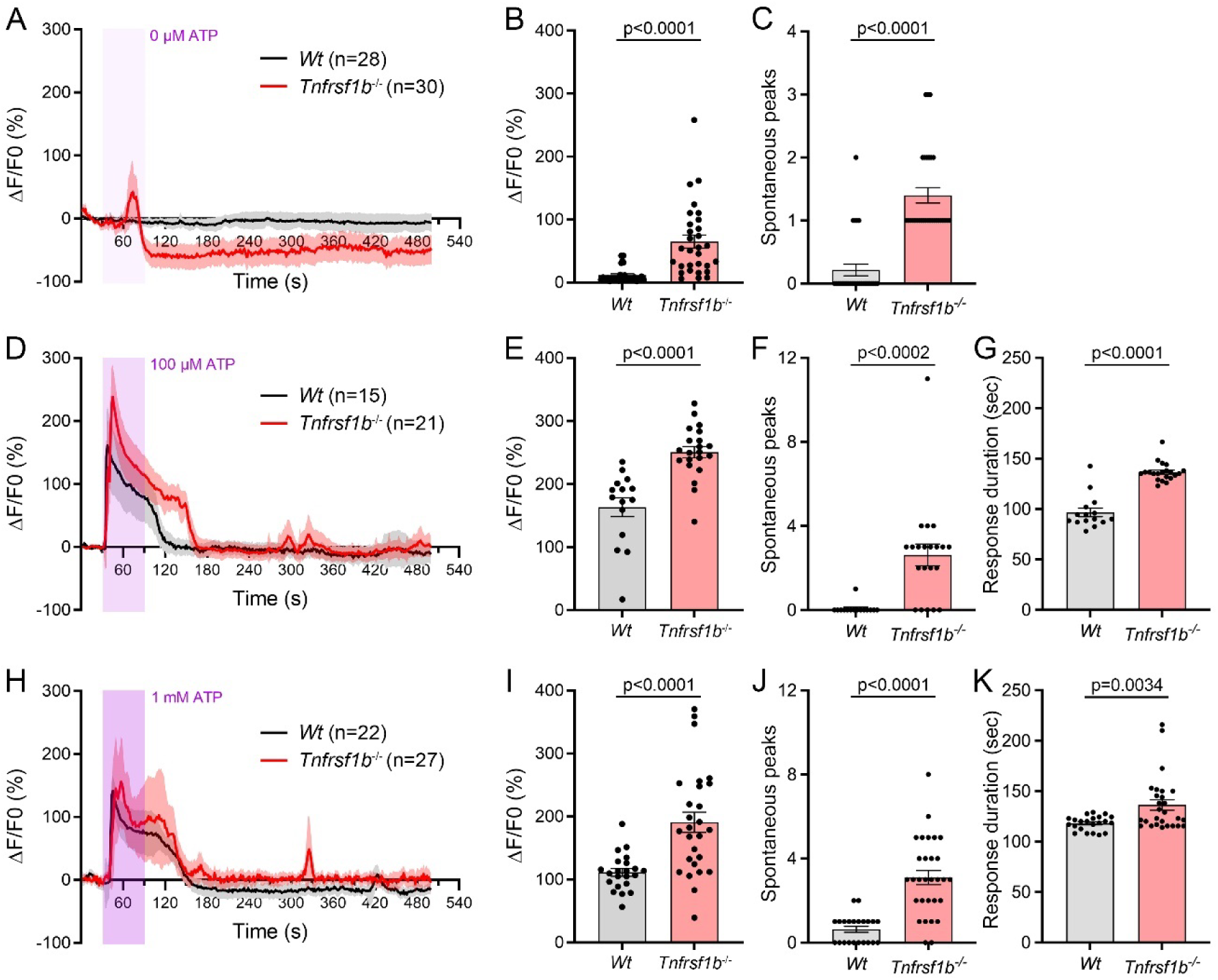
TNFR2 regulates astrocyte Ca^2+^ transients. (A, D, H) [Ca^2+^]_i_ transients in *Wt* and *Tnfrsf1b^-/-^* primary astrocytes after exposures to vehicle or ATP and measured by % increase of Fluo4 fluorescence above baseline (ΔF/F). Shaded purple areas represent vehicle or ATP perfusion intervals. (B, E, I) Peak % of Fluo4 ΔF/F after exposure to vehicle or ATP; (C, F, J) Number of spontaneous [Ca^2+^]_i_ peaks after exposure to vehicle or ATP. (G, K) Duration of the [Ca^2+^]_i_ response to ATP. Each point represents a recorded cell. Cells were recorded from 3 replicates/condition. Results represent average ± SEM; Groups compared by Student’s *t* test with p≤0.05 considered statistically significant.

### 3.9. Ablation of TNFR2 alters the gene expression profile of hippocampal astrocytes in physiological conditions

To identify pathways/genes regulated by TNFR2 in astrocytes that could be directly or indirectly responsible for modulating hippocampal synaptic function in physiological conditions, we performed bulk RNAseq of hippocampal astrocytes isolated from adult *Gfap^creERT2^:Tnfrsf1b^fl/fl^* and *Tnfrsf1b^fl/fl^* male mice. We identified 164 differentially expressed genes (DEGs), with 13% (22 genes) upregulated and 87% (142 genes) downregulated in *Gfap^creERT2^:Tnfrsf1b^fl/fl^* mice compared to controls (Fig. 10A). Notably, among the top 50 most significant DEGs, only 1 was upregulated, while the remaining 49 were downregulated (Fig. 10B) (GEO:GSE288931).

**Figure 10.**
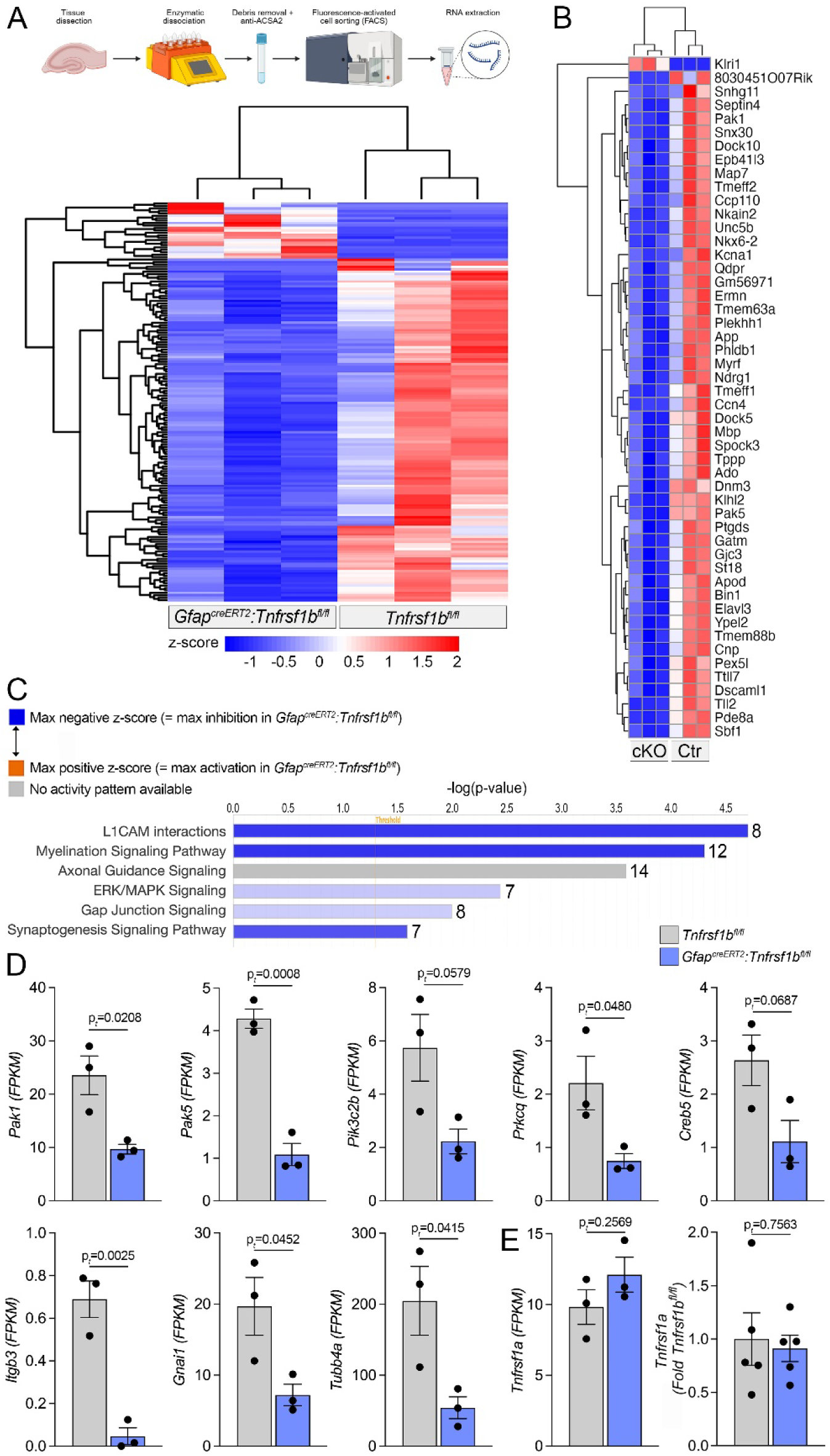
TNFR2 ablation alters gene expression in hippocampal astrocytes in physiological conditions. (A, B)) Heatmap of all 164 (A) and of the top 50 (B) differentially expressed genes (*p*adj≤0.05) in hippocampal astrocytes from *Gfap^creERT2^:Tnfrsf1b^fl/fl^* and *Tnfrsf1b^fl/fl^* male mice; in blue = downregulated genes; in red = upregulated genes. (C) Key affected canonical pathways identified by Ingenuity Pathway Analysis (IPA), where DEGs with an absolute log2 fold change (|LogFc|) ≥ 1.0, and a *p*adj≤0.1 were considered statistically significant. The number of differentially expressed genes in each pathway is depicted on the right of the bar graph. (D) Expression profiles of select genes identified across multiple pathways altered between *Gfap^creERT2^:Tnfrsf1b^fl/fl^* and *Tnfrsf1b^fl/fl^* astrocyte expression levels are shown as Fragments Per Kilobase per million Mapped reads (FPKM). (E) *Tnfrsf1a* expression by RNAseq and qPCR; n=3-5/group, p values after Student’s *t* test are indicated on each graph. Results represent average ± SEM.

To better understand the biological relevance of these DEGs, we performed ingenuity pathway analysis (IPA). This revealed numerous canonical pathways significantly altered in hippocampal astrocytes lacking TNFR2, with all but one pathway being downregulated (Fig. S1). Key affected pathways included those involved in adhesion molecule interactions, myelination, axonal guidance, ERK/MAPK signaling, gap junctions, and synaptogenesis, all of which were downregulated in *Gfap^creERT2^:Tnfrsf1b^fl/fl^* mice compared to *Tnfrsf1b^fl/fl^* controls (Fig. 10C). These pathways play essential roles in maintaining physiological synaptic function, emphasizing the importance of astrocytic TNFR2 in this context. Furthermore, these pathways were found to be interconnected through shared molecules, with genes such as *Pak1*, *Pak5*, *PiK3c2b*, *Prkcq*, *Creb5*, *Itgb3*, *Gnai*, and *Tubb4a* common to several of them (Fig. 10D). These findings emphasize the intricate regulatory role of astroglial TNFR2, on which multiple pathways critical for hippocampal synaptic function and neuronal communication seem to converge.

Furthermore, we excluded that the gene expression changes associated with astroglial TNFR2 ablation were dependent on adaptive dysregulation of the TNFR1 receptor, since both by RNAseq analysis and by qPCR quantification in sorted hippocampal astrocytes we found no differences between *Gfap^creERT2^:Tnfrsf1b^fl/fl^* mice and *Tnfrsf1b^fl/fl^* controls (Fig. 10E).

## 4. Discussion

In the present work, we demonstrate that TNFR2 signaling in astrocytes plays an essential role in regulating hippocampal synaptic function, plasticity, and cognition in physiological conditions. This is the first report to implicate TNFR2 in these processes. Furthermore, we uncover sex-specific differences in TNFR2 functions, with males relying more on astroglial TNFR2 than females for physiological synaptic function, as shown by the exacerbated synaptic and cognitive disturbances manifested by male mice lacking TNFR2 compared to females.

The rationale driving our study is twofold. First, the gap of knowledge in our understanding of astroglial TNFR2 function in synaptic modulation: indeed, the established role of TNF in astrocyte physiology and, as a result, in the regulation of synaptic and cognitive function has been exclusively associated to TNFR1 [2, 30, 59]. No studies have queried how astrocyte physiology might be regulated by TNFR2, possibly due to its minimal expression levels in astrocytes under physiological conditions [60, 61]. Second, the body of evidence demonstrating important protective roles for glial TNFR2 in both physiological and pathological conditions: indeed, TNFR2 in microglia, mature oligodendrocytes and oligodendrocyte precursor cells (OPCs) has been shown to drive cell survival, anti-inflammatory and immunomodulatory effects that positively support cell homeostasis as well as the cellular response to disease/damage [34, 40, 41, 46, 62], suggesting that TNFR2 might have similar functions in astrocytes as well.

To investigate the role of astroglial TNFR2, we generated *Gfap^creERT2^:Tnfrsf1b^fl/fl^* mice with selective ablation of TNFR2 in astrocytes and probed for TNFR2 dependent effects in the hippocampus, the key center for higher order cognition, learning and memory. Compared to *Tnfrsf1b^fl/fl^* control mice, *Gfap^creERT2^:Tnfrsf1b^fl/fl^* mice exhibited dysregulation of a broad spectrum of proteins necessary for synaptic function. This was most evident in male mice where pre-synaptic SNARE proteins SNAP-25 and Synaptotagmin 1/2, post-synaptic proteins PSD-93 and PSD-95, and glutamate receptor subunits were all found to be upregulated. As these proteins are expressed predominantly in neurons, their increased expression suggests that ablating TNFR2 from astrocytes impacts hippocampal neuronal function, indicating that astroglial TNFR2 is needed for proper communication between astrocytes and neurons.

SNAP-25, which is essential for vesicle fusion, is also involved in the regulation of intracellular calcium dynamics, whereby its phosphorylation inhibits neuronal voltage-gated calcium channels (VGCCs) [63]. Overexpression of SNAP-25 in hippocampal neurons significantly reduces calcium responses to depolarization by inhibiting VGCCs [64]. Therefore, it is possible that upregulation of SNAP-25 following astroglial TNFR2 ablation may contribute to the impairment of physiological synaptic transmission.

Post-synaptic proteins PSD-93 and PSD-95 have critical roles in AMPA receptor trafficking and in forming NMDA receptor-associated signaling complexes involved in synaptic plasticity [65]. As such, their upregulation may lead to alterations of homeostatic synaptic function. Indeed, increased PSD-95 has been shown to inhibit synaptic scaling [66], reducing the strength of synaptic connections during periods of excessive activity. Upregulation of PSD-95 and PSD-93 due to astroglial TNFR2 ablation may lead to inhibition of synaptic scaling, potentially leading to excessive excitatory neurotransmission and excitotoxicity.

Male *Gfap^creERT2^:Tnfrsf1b^fl/fl^* mice also exhibited altered expression of AMPA receptors (AMPARs) and NMDA receptors (NMDARs) subunits. As AMPARs facilitate the rapid strengthening of synaptic connections [67], and NMDARs trigger the calcium-dependent signaling required for long-term potentiation (LTP) [68], both are crucial for synaptic plasticity. The number and subunit composition of AMPARs determine the efficiency and dynamics of AMPAR-mediated synaptic signaling [69]. Mice lacking astroglial TNFR2 showed upregulation of GluA1 and GluA2/3 AMPAR subunits. The GluA2 subunit is particularly important as it controls the receptor’s permeability to Ca^2+^, in that GluA2 containing AMPARs are impermeable to calcium [70]. Consequently, upregulation of GluA2 as seen with astroglial TNFR2 ablation may negatively impact synaptic plasticity.

With respect to NMDARs, we tested the expression of the two obligatory subunits GluN1, which contains the glycine/D-serine-binding site, and GluN2, which contains the binding site for glutamate [71]. GluN1 did not change between *Gfap^creERT2^:Tnfrsf1b^fl/fl^* and *Tnfrsf1b^fl/fl^* mice, but GluN2A, which with GluN2B constitutes the subtype most highly expressed in the hippocampus [71], was upregulated in mice lacking astroglial TNFR2. This is significant since GluN2A is enriched in NMDARs located at synaptic sites where they play a crucial role in synaptic plasticity [72]. Thus, upregulation of GluN2A upon astroglial TNFR2 ablation may also contribute to an impairment in synaptic functionality. Notably, these same proteins where minimally affected in female *Gfap^creERT2^:Tnfrsf1b^fl/fl^* mice, with only mild downregulation of GluA2/3, contrary to male mice, and a mild upregulation of GluN2A, in line with male mice. Interestingly, in both male and female *Gfap^creERT2^:Tnfrsf1b^fl/fl^* mice the glutamate transporter EAAT1, primarily expressed in astrocytes, was significantly upregulated compared to controls. As astrocytic EAAT1 reuptakes and clears glutamate from the extracellular space to maintain glutamate homeostasis [73], this finding suggests that astroglial TNFR2 exerts a direct effect on synaptic transmission by controlling the availability of glutamate at the synaptic cleft [74]. We speculate that the upregulation of EAAT1 may occur as a feedback response of astrocytes to the upregulation of neuronal-specific synaptic proteins and glutamate receptors that cause enhanced glutamatergic signaling. The increased EAAT1 levels would enhance glutamate reuptake to mitigate potential excitotoxic damage. Furthermore, our data underscore that TNFR2 in astrocytes participates in the regulation of homeostatic synaptic function both via astroglial (direct) and neuronal (indirect) mechanisms, thus serving as a complete controller of neuron-glia communication.

One potential mechanism through which astroglial TNFR2 may participate in synaptic regulation is by controlling cell reactivity, in that its steady state activation is needed to maintain astrocytes, and indirectly neurons, in the optimal homeostatic conditions to execute proper synaptic transmission. As such, we hypothesized that lack of TNFR2 directed astrocytes to acquire an activated phenotype, as they would in response to stress signals like excessive glutamate or ATP released from damaged neural cells. In these conditions, astrocytes become reactive, displaying changes like increased branching and process elongation, and increased proliferation [54]. To this end, we found that in male mice lacking astroglial TNFR2 hippocampal astrocytes were both hypertrophic and hyperplasic to indicate reactivity and gliosis. Since this phenomenon is typically accompanied by transcriptional changes that lead to the secretion of immunomodulatory and inflammatory factors that affect neighboring cells such as microglia, we evaluated microglia numbers and morphology in the same hippocampal regions. As with astrocytes, microglia numbers were elevated in mice lacking astroglial TNFR2 indicating proliferation, but no morphological changes were observed. This enhanced glia reactivity was confirmed with our TSPO binding studies that showed global increase of glial reactivity in the brain, and specifically in hippocampus, cortex and thalamus in male *Gfap^creERT2^:Tnfrsf1b^fl/fl^* mice. With the assays we performed, we could not identify any specific cytokine potentially released by reactive TNFR2 deficient astrocytes that might contribute to this phenomenon. Yet, this does not exclude that astroglial TNFR2 signaling might maintain homeostatic synaptic function via modulation (specifically suppression) of detrimental inflammatory activation. Indeed, such a function has been already demonstrated in previous studies from our lab for microglial and oligodendroglial TNFR2 [40, 41, 46, 62, 75]. Further studies are warranted to fully address this question.

The increased number of neurons in the dentate gyrus of male mice lacking astroglial TNFR2 suggests that TNFR2 in astrocytes may influence adult neurogenesis. Astrocytes in particular, have been shown to control the growth and differentiation of adult-born granule cells via release of trophic factors and gliotransmitters [76], such as brain derived neurotrophic factor (BDNF) and fibroblast growth factor 2 (FGF2) [77, 78]. Furthermore, administration of D-serine was shown to increase adult hippocampal neurogenesis by increasing proliferation and survival of newborn neurons [79]. We speculate that the reactive state of hippocampal astrocytes following astroglial TNFR2 ablation may alter the release of gliotransmitters and other soluble factors that influence neurogenesis, resulting in the increase in neurons we observed in the dentate gyrus. Further studies are needed to explore this possibility.

The profound changes observed in the hippocampus of male mice led us to postulate these might translate in functional deficits, including impaired synaptic plasticity and cognition, both of which were confirmed by the suppressed LTP and the impairment of learning and memory assessed in both male and female *Gfap^creERT2^:Tnfrsf1b^fl/fl^* mice. A potential explanation for this effect is the saturation of synaptic plasticity mechanisms due to increased AMPAR-mediated synaptic strength [80]. Indeed, incorporation of AMPARs into synapses is a well-established mechanism for the expression of LTP [81], and previous studies have shown that glial TNF increases synaptic strength by increasing the surface expression of AMPARs [28]. Furthermore, it has been shown that higher surface levels of GluA1-containing AMPARs alter AMPA receptor trafficking, leading to suppressed LTP at hippocampal synapses [80]. As we see upregulation of the GluA1 and suppressed LTP in *Gfap^creERT2^:Tnfrsf1b^fl/fl^* male mice, this may represent a mechanism by which astroglial TNFR2 regulates hippocampal synaptic plasticity.

To address how TNFR2 in astrocytes may trigger the cascade of events modulating synaptic function, we probed for intracellular calcium dynamics, the primary mode of astrocyte communication. Indeed, fluctuations in intracellular calcium are the critical signal for gliotransmitter release [82]. We found that TNFR2 ablated astrocytes had aberrantly elevated calcium transients in both non-stimulated and stimulated conditions. We speculate that this might lead to increased release of gliotransmitters such as glutamate and D-serine, which in turn would activate presynaptic NMDA receptors and increase excitatory transmission. Further studies are warranted to validate this hypothesis.

To gain insight into the mechanisms of astroglial TNFR2-dependent modulation of physiological hippocampal synaptic plasticity, we performed RNAseq analysis of the hippocampal astrocyte transcriptome of naïve *Gfap^creERT2^:Tnfrsf1b^fl/fl^* male mice compared to controls. The overall picture that emerged depicts TNFR2 as essential for astrocytic regulation of synaptic function. Among the most significantly dysregulated pathways identified by ingenuity pathway analysis (IPA) is the one centered on L1 cell adhesion molecule (L1CAM) interactions. Cell adhesion molecules are crucial for cell adhesion, migration, and signaling, and L1-CAMs have been shown to play key roles in neurogenesis, neural plasticity, and learning and memory in the adult nervous system [83–86]. In addition, synaptogenesis and gap junction signaling were also among the top dysregulated pathways. It is well established that astrocytes regulate glutamatergic synaptogenesis by secreting various factors [87], and this is one of the mechanisms by which astrocytes modulate synaptic plasticity. Interestingly, the dysregulated pathways we identified were interconnected by a core group of shared downregulated genes, *Pak1*, *Pik3c2b*, *Creb5*, and *Tubb4a*, pointing at these molecules as key signals in TNFR2-dependent regulation of astrocyte-neuron communication. Noteworthy is that axonal guidance signaling was also found to be disrupted, with 14 differentially regulated genes, although no specific up or down pattern could be identified. In the adult nervous system, canonical axonal guidance proteins such as semaphorins and ephrins have been shown to function as regulators of synaptic plasticity [88, 89]. In our model, hippocampal astrocytes lacking TNFR2 exhibited downregulation of ephrin type-A receptor 3 (*Epha3*), semaphorin-6A (*Sema6a*), and semaphorin receptor 5A (*Plxnb3*). Epha3 has been shown to be highly expressed in the adult mouse hippocampus, where it is localized on astrocytic processes that envelop dendritic spines [90]. Astrocytic Epha3 interacts with ligand EphA4 on dendritic spines of pyramidal neurons to regulate structural formation and excitatory connection [90]. Despite not observing any significant alterations in dendritic spine morphology with ablation of astroglial TNFR2, it is possible this downregulation of astrocytic *Epha3* due to lack of TNFR2 plays a role in altering synaptic transmission and plasticity. With respect to Sema6a, an *in vitro* study showed that astrocytic Sema6a acts to repel OPCs from blood vessels allowing their differentiation into mature oligodendrocytes, which represents a mechanism by which astrocytes regulate OPC migration and differentiation [91]. Since *Sema6a* is downregulated in astrocytes lacking TNFR2, this could mean that TNFR2 signaling in astrocytes impact OPCs and oligodendrocyte function leading to altered myelination, thus synaptic transmission. Indeed, astrocytes have been shown to promote myelination by various mechanisms, including secreting factors that influence proliferation, differentiation, and migration of oligodendrocytes [92], and there is evidence that TNFR2 expressed in astrocytes via secretion of CXCL12 promotes OPC differentiation and myelin formation [32]. Thus, since we found that one of the top differentially expressed pathways in astrocytes lacking TNFR2 is *myelination signaling*, it is possible that astroglial TNFR2 signaling affects synaptic plasticity via regulation of oligodendrocyte functions, including myelination.

It is important to underscore that the effects observed in male mice were only minimally recapitulated in female mice, adding to the growing body of evidence that there are deep differences in the regulation of neural functions between females and males, both in health and disease. It is well known that sex hormones influence CNS functions [93], including synaptic plasticity and long-term memory [94], with several studies demonstrating that estradiol improves synaptic plasticity and memory [95–98]. Estrogens also exert neuroprotective effects such as reducing proliferation and activation of astrocytes in pathological conditions [99]. These regulatory and protective effects of estrogen may contribute to the reduced detrimental effects we observed in female mice lacking astroglial TNFR2. Furthermore, the rodent estrous cycle has been found to influence a variety of physiological mechanisms such as cell proliferation, hippocampal volume, and LTP [100–102]. As we did not account for the estrous cycle phases in our study, it is possible that fluctuations in hormone levels may have contributed to the sex differences observed in our findings.

## 5. Conclusions

In conclusion, we show that lack of TNFR2 in astrocytes leads to: altered expression of key proteins involved in hippocampal synaptic transmission and plasticity, increased hippocampal gliosis, impaired LTP, and deficits in learning and memory. These effects were associated with dysregulation of astroglial calcium dynamics and of genes associated with astrocyte interactions and communication with neighboring cells, especially neurons and oligodendroglia. Such effects were more prominent in male mice. Together, our findings demonstrate that astroglial TNFR2 is an essential player in the maintenance of homeostatic synaptic function, plasticity and cognition, especially in males. While this adds new knowledge on the role of TNFR2 in the CNS, it also warrants more studies to further delve into the associated molecular mechanisms, including the possible determinants of sex bias, as well as to probe for TNFR2 function in the context of CNS disease and trauma, particularly if associated with cognitive impairment.

## Author contributions

BNC and PI performed most of the experiments and conducted data analysis; BNC wrote the manuscript; TMP performed immunohistochemistry, imaging, Sholl analysis on astrocytes, and data analysis; HLD assisted with FACS and RNAseq data analysis; SM and AIA performed immunohistochemistry, imaging, stereological counting; AIA analyzed dendritic spine morphology and performed behavioral assays; SJ conducted Sholl analysis on neurons; MCA performed Golgi-Cox staining and imaging; PKS performed RNAseq data analysis; DJT and CMA performed LTP experiments and data analysis; BAP and JDM performed [^3^H]PBR28 autoradiography and data analysis; LW and LB performed calcium imaging experiments and data analysis; KLL performed multiplex analysis; RB conceived the study, conducted data analysis and interpretation, and wrote the manuscript.

## Funding

This work was supported by: NIH NINDS grants 1R01NS094522-01 and 1R21NS120028-01 (RB); Italian Multiple Sclerosis Foundation (FISM) grant 2020/R-Single/024 (RB); Team Science Award Program, Office of the Executive Dean For Research, University of Miami Miller School of Medicine (RB); The Miami Project to Cure Paralysis and the Buoniconti Fund (RB); NIH NINDS grants R01NS127146, R01NS105616, R21NS137641 (LB); NIH NIA grant R21AG087451 (LB); Novo Nordisk Foundation grant NNF22OC0079804 (KLL).

## Acknowledgements

We thank Eva Juarez, Shaffiat Karmally, Estrid Thougaard, Antonella Mini, Nitya Anne and Henri Chédotal for their assistance with animal colony maintenance, genotyping and tissue processing. We thank Ulla Munk for assistance with multiplex analysis, and Dr. Oliver Umland for assistance with FACS. Graphical abstract and schematic in Fig. 10A were generated with BioRender.com. The accession number for the sequencing data reported in this paper is GEO:GSE288931.

## Abbreviations

aCSF: Artificial cerebral spinal fluid
BDNF: Brain derived neurotrophic factor
BSA: Bovine serum albumin
DG: Dentate gyrus
DMEM: Dulbecco’s modified Eagle’s medium
EYFP: Enhanced yellow fluorescent protein
FACS: Fluorescence activated cell sorting
FBS: Fetal bovine serum
FC: Fear conditioning
FGF2: Fibroblast growth factor 2
fEPSP: Field excitatory postsynaptic potentials
GFAP: Glial fibrillary acidic protein
GFP: Green fluorescent protein
I/O: Input/output
IPA: Ingenuity pathway analysis
LTP: Long-term potentiation
MAGUKs: Membrane-associated guanylate kinases
MWM: Morris water maze
NOR: Novel Object Recognition
OD: Optical density
PET: Positron emission tomography
PPF: Paired-pulse facilitation
ROI: Region of interest
solTNF: Soluble tumor necrosis factor
tmTNF: Transmembrane tumor necrosis factor
TNF: tumor necrosis factor
TNFR1: Tumor necrosis factor receptor 1
TNFR2: Tumor necrosis factor receptor 2
TSPO: Translocator protein 18 kDa
VGCCs: Voltage-gated calcium channels
WT: Wild type

## Notes

### Competing Interest Statement

The authors have declared no competing interest.

## References

1. Turrigiano, G.G. and S.B. Nelson, Homeostatic plasticity in the developing nervous system. Nat Rev Neurosci, 2004. 5(2): p. 97–107.

2. Santello, M., N. Toni, and A. Volterra, Astrocyte function from information processing to cognition and cognitive impairment. Nat Neurosci, 2019. 22(2): p. 154–166.

3. Araque, A., et al., Tripartite synapses: glia, the unacknowledged partner. Trends Neurosci, 1999. 22(5): p. 208–15.

4. Perea, G., M. Navarrete, and A. Araque, Tripartite synapses: astrocytes process and control synaptic information. Trends Neurosci, 2009. 32(8): p. 421–31.

5. Derouiche, A., et al., Anatomical aspects of glia-synapse interaction: the perisynaptic glial sheath consists of a specialized astrocyte compartment. J Physiol Paris, 2002. 96(3-4): p. 177–82.

6. Reichenbach, A., A. Derouiche, and F. Kirchhoff, Morphology and dynamics of perisynaptic glia. Brain Res Rev, 2010. 63(1-2): p. 11–25.

7. Ota, Y., A.T. Zanetti, and R.M. Hallock, The role of astrocytes in the regulation of synaptic plasticity and memory formation. Neural Plast, 2013. 2013: p. 185463.

8. Araque, A., et al., Gliotransmitters travel in time and space. Neuron, 2014. 81(4): p. 728–39.

9. Oliet, S.H., R. Piet, and D.A. Poulain, Control of glutamate clearance and synaptic efficacy by glial coverage of neurons. Science, 2001. 292(5518): p. 923–6.

10. Allen, N.J. and B.A. Barres, Signaling between glia and neurons: focus on synaptic plasticity. Curr Opin Neurobiol, 2005. 15(5): p. 542–8.

11. Chung, W.S., et al., Astrocytes mediate synapse elimination through MEGF10 and MERTK pathways. Nature, 2013. 504(7480): p. 394–400.

12. Chung, W.S., N.J. Allen, and C. Eroglu, Astrocytes Control Synapse Formation, Function, and Elimination. Cold Spring Harb Perspect Biol, 2015. 7(9): p. a020370.

13. Christopherson, K.S., et al., Thrombospondins are astrocyte-secreted proteins that promote CNS synaptogenesis. Cell, 2005. 120(3): p. 421–33.

14. Maggio, N.V., A., Tumor necrosis factor (TNF) modulates synaptic plasticity in a concentrationdependent manner through intracellular calcium stores. 2018, J Mol Med (Berl). p. 1039–1047.

15. Black, R.A., et al., A metalloproteinase disintegrin that releases tumour-necrosis factor-alpha from cells. Nature, 1997. 385(6618): p. 729–33.

16. Kalliolias, G.D. and L.B. Ivashkiv, TNF biology, pathogenic mechanisms and emerging therapeutic strategies. Nat Rev Rheumatol, 2016. 12(1): p. 49–62.

17. Grell, M., et al., The type 1 receptor (CD120a) is the high-affinity receptor for soluble tumor necrosis factor. Proc Natl Acad Sci U S A, 1998. 95(2): p. 570–5.

18. Henneberger, C., et al., Long-term potentiation depends on release of D-serine from astrocytes. Nature, 2010. 463(7278): p. 232–6.

19. Brambilla, R., et al., Inhibition of soluble tumour necrosis factor is therapeutic in experimental autoimmune encephalomyelitis and promotes axon preservation and remyelination. Brain, 2011. 134(Pt 9): p. 2736–54.

20. Kim, J.E., et al., Tumor necrosis factor-alpha-mediated threonine 435 phosphorylation of p65 nuclear factor-kappaB subunit in endothelial cells induces vasogenic edema and neutrophil infiltration in the rat piriform cortex following status epilepticus. J Neuroinflammation, 2012. 9: p. 6.

21. Magliozzi, R., et al., Meningeal inflammation changes the balance of TNF signalling in cortical grey matter in multiple sclerosis. J Neuroinflammation, 2019. 16(1): p. 259.

22. Heir, R., et al., Astrocytes Are the Source of TNF Mediating Homeostatic Synaptic Plasticity. J Neurosci, 2024. 44(14).

23. Brambilla, R., The contribution of astrocytes to the neuroinflammatory response in multiple sclerosis and experimental autoimmune encephalomyelitis. Acta Neuropathol, 2019.

24. Di Castro, M.A. and A. Volterra, Astrocyte control of the entorhinal cortex-dentate gyrus circuit: Relevance to cognitive processing and impairment in pathology. Glia, 2022. 70(8): p. 1536–1553.

25. Stellwagen, D. and R.C. Malenka, Synaptic scaling mediated by glial TNF-alpha. Nature, 2006. 440(7087): p. 1054–9.

26. Domercq, M., et al., P2Y1 receptor-evoked glutamate exocytosis from astrocytes: control by tumor necrosis factor-alpha and prostaglandins. J Biol Chem, 2006. 281(41): p. 30684–96.

27. Bezzi, P., et al., CXCR4-activated astrocyte glutamate release via TNFalpha: amplification by microglia triggers neurotoxicity. Nat Neurosci, 2001. 4(7): p. 702–10.

28. Beattie, E.C., et al., Control of synaptic strength by glial TNFalpha. Science, 2002. 295(5563): p. 2282–5.

29. Pribiag, H. and D. Stellwagen, TNF-α downregulates inhibitory neurotransmission through protein phosphatase 1-dependent trafficking of GABA(A) receptors. J Neurosci, 2013. 33(40): p. 15879–93.

30. Habbas, S., et al., Neuroinflammatory TNFalpha Impairs Memory via Astrocyte Signaling. Cell, 2015. 163(7): p. 1730–41.

31. Veroni, C., et al., Connecting Immune Cell Infiltration to the Multitasking Microglia Response and TNF Receptor 2 Induction in the Multiple Sclerosis Brain. Front Cell Neurosci, 2020. 14: p. 190.

32. Patel, J.R., et al., Astrocyte TNFR2 is required for CXCL12-mediated regulation of oligodendrocyte progenitor proliferation and differentiation within the adult CNS. Acta Neuropathol, 2012. 124(6): p. 847–60.

33. Casper, K.B., K. Jones, and K.D. McCarthy, Characterization of astrocyte-specific conditional knockouts. Genesis, 2007. 45(5): p. 292–9.

34. Madsen, P.M., et al., Oligodendroglial TNFR2 Mediates Membrane TNF-Dependent Repair in Experimental Autoimmune Encephalomyelitis by Promoting Oligodendrocyte Differentiation and Remyelination. J Neurosci, 2016. 36(18): p. 5128–43.

35. Noldus, L.P.J.J., A.J. Spink, and R.A.J. Tegelenbosch, EthoVision: A versatile video tracking system for automation of behavioral experiments. Behavior Research Methods Instruments & Computers, 2001. 33(3): p. 398–414.

36. Leger, M., et al., Object recognition test in mice. Nature Protocols, 2013. 8(12): p. 2531–2537.

37. Tapanes, S.A., et al., Inhibition of glial D-serine release rescues synaptic damage after brain injury. Glia, 2022. 70(6): p. 1133–1152.

38. Titus, D.J., et al., A negative allosteric modulator of PDE4D enhances learning after traumatic brain injury. Neurobiology of Learning and Memory, 2018. 148: p. 38–49.

39. Takao, K. and T. Miyakawa, Light/dark transition test for mice. J Vis Exp, 2006(1): p. 104.

40. Gao, H., et al., Opposing Functions of Microglial and Macrophagic TNFR2 in the Pathogenesis of Experimental Autoimmune Encephalomyelitis. Cell Rep, 2017. 18(1): p. 198–212.

41. Madsen, P.M., et al., Oligodendrocytes modulate the immune-inflammatory response in EAE via TNFR2 signaling. Brain Behav Immun, 2020. 84: p. 132–146.

42. Mikkelsen, J.D., et al., Characterization of the Novel P2X7 Receptor Radioligand [ACS Chem Neurosci, 2023. 14(1): p. 111–118.

43. Thougaard, E., et al., Systemic treatment with a selective TNFR2 agonist alters the central and peripheral immune responses and transiently improves functional outcome after experimental ischemic stroke. J Neuroimmunol, 2023. 385: p. 578246.

44. Risher, W.C., et al., Rapid Golgi analysis method for efficient and unbiased classification of dendritic spines. PLoS One, 2014. 9(9): p. e107591.

45. Titus, D.J., et al., Age-dependent alterations in cAMP signaling contribute to synaptic plasticity deficits following traumatic brain injury. Neuroscience, 2013. 231: p. 182–94.

46. Desu, H.L., et al., TNFR2 Signaling Regulates the Immunomodulatory Function of Oligodendrocyte Precursor Cells. Cells, 2021. 10(7).

47. Desu, H.L., et al., TNFR2 signaling in oligodendrocyte precursor cells suppresses their immune-inflammatory function and detrimental microglia activation in CNS demyelinating disease. Brain Behav Immun, 2024. 123: p. 81–98.

48. Jahn, R., D.C. Cafiso, and L.K. Tamm, Mechanisms of SNARE proteins in membrane fusion. Nat Rev Mol Cell Biol, 2024. 25(2): p. 101–118.

49. Owen, D. R.e.a., wo binding sites for [3H]PBR28 in human brain: implications for TSPO PET imaging of neuroinflammation. 2010: J Cereb Blood Flow Metab.

50. Lavisse, S., et al., Reactive astrocytes overexpress TSPO and are detected by TSPO positron emission tomography imaging. J Neurosci, 2012. 32(32): p. 10809–18.

51. Gouilly, D., et al., Neuroinflammation PET imaging of the translocator protein (TSPO) in Alzheimer’s disease: An update. Eur J Neurosci, 2022. 55(5): p. 1322–1343.

52. Porro, C., A. Cianciulli, and M.A. Panaro, The Regulatory Role of IL-10 in Neurodegenerative Diseases. Biomolecules, 2020. 10(7).

53. Woodburn, S.C., J.L. Bollinger, and E.S. Wohleb, The semantics of microglia activation: neuroinflammation, homeostasis, and stress. J Neuroinflammation, 2021. 18(1): p. 258.

54. Brambilla, R., The contribution of astrocytes to the neuroinflammatory response in multiple sclerosis and experimental autoimmune encephalomyelitis. Acta Neuropathol, 2019. 137(5): p. 757–783.

55. Selden, N.R., et al., Complementary roles for the amygdala and hippocampus in aversive conditioning to explicit and contextual cues. Neuroscience, 1991. 42(2): p. 335–50.

56. Bazargani, N. and D. Attwell, Astrocyte calcium signaling: the third wave. Nat Neurosci, 2016. 19(2): p. 182–9.

57. Parpura, V., et al., Glutamate-mediated astrocyte-neuron signalling. Nature, 1994. 369(6483): p. 744–7.

58. Pasti, L., et al., Intracellular calcium oscillations in astrocytes: a highly plastic, bidirectional form of communication between neurons and astrocytes in situ. J Neurosci, 1997. 17(20): p. 7817–30.

59. Santello, M., P. Bezzi, and A. Volterra, TNFalpha controls glutamatergic gliotransmission in the hippocampal dentate gyrus. Neuron, 2011. 69(5): p. 988–1001.

60. Zhang, Y., et al., An RNA-sequencing transcriptome and splicing database of glia, neurons, and vascular cells of the cerebral cortex. J Neurosci, 2014. 34(36): p. 11929–47.

61. Zhang, Y., et al., Purification and Characterization of Progenitor and Mature Human Astrocytes Reveals Transcriptional and Functional Differences with Mouse. Neuron, 2016. 89(1): p. 37–53.

62. Desu, H.L., et al., TNFR2 signaling in oligodendrocyte precursor cells suppresses their immune-inflammatory function and detrimental microglia activation in CNS demyelinating disease. Brain Behav Immun, 2024. 123: p. 81–98.

63. Pozzi, D., et al., Activity-dependent phosphorylation of Ser187 is required for SNAP-25-negative modulation of neuronal voltage-gated calcium channels. Proc Natl Acad Sci U S A, 2008. 105(1): p. 323–8.

64. Verderio, C., et al., SNAP-25 modulation of calcium dynamics underlies differences in GABAergic and glutamatergic responsiveness to depolarization. Neuron, 2004. 41(4): p. 599–610.

65. Carlisle, H.J., et al., Opposing effects of PSD-93 and PSD-95 on long-term potentiation and spike timing-dependent plasticity. J Physiol, 2008. 586(24): p. 5885–900.

66. Sun, Q. and G.G. Turrigiano, PSD-95 and PSD-93 play critical but distinct roles in synaptic scaling up and down. J Neurosci, 2011. 31(18): p. 6800–8.

67. Diering, G.H. and R.L. Huganir, The AMPA Receptor Code of Synaptic Plasticity. Neuron, 2018. 100(2): p. 314–329.

68. Morris, R.G., et al., Selective impairment of learning and blockade of long-term potentiation by an N-methyl-D-aspartate receptor antagonist, AP5. Nature, 1986. 319(6056): p. 774–6.

69. Gan, Q., C.L. Salussolia, and L.P. Wollmuth, Assembly of AMPA receptors: mechanisms and regulation. J Physiol, 2015. 593(1): p. 39–48.

70. Isaac, J.T., M.C. Ashby, and C.J. McBain, The role of the GluR2 subunit in AMPA receptor function and synaptic plasticity. Neuron, 2007. 54(6): p. 859–71.

71. Shipton, O.A. and O. Paulsen, GluN2A and GluN2B subunit-containing NMDA receptors in hippocampal plasticity. Philos Trans R Soc Lond B Biol Sci, 2014. 369(1633): p. 20130163.

72. Paoletti, P., C. Bellone, and Q. Zhou, NMDA receptor subunit diversity: impact on receptor properties, synaptic plasticity and disease. Nat Rev Neurosci, 2013. 14(6): p. 383–400.

73. Mahmoud, S., et al., Astrocytes Maintain Glutamate Homeostasis in the CNS by Controlling the Balance between Glutamate Uptake and Release. Cells, 2019. 8(2).

74. Malik, A.R. and T.E. Willnow, Excitatory Amino Acid Transporters in Physiology and Disorders of the Central Nervous System. Int J Mol Sci, 2019. 20(22).

75. Madsen, P.M., et al., Mitochondrial DNA Double-Strand Breaks in Oligodendrocytes Cause Demyelination, Axonal Injury, and CNS Inflammation. J Neurosci, 2017. 37(42): p. 10185–10199.

76. Sultan, S., et al., Synaptic Integration of Adult-Born Hippocampal Neurons Is Locally Controlled by Astrocytes. Neuron, 2015. 88(5): p. 957–972.

77. Quesseveur, G., et al., BDNF overexpression in mouse hippocampal astrocytes promotes local neurogenesis and elicits anxiolytic-like activities. Transl Psychiatry, 2013. 3(4): p. e253.

78. Rai, K.S., B. Hattiangady, and A.K. Shetty, Enhanced production and dendritic growth of new dentate granule cells in the middle-aged hippocampus following intracerebroventricular FGF-2 infusions. Eur J Neurosci, 2007. 26(7): p. 1765–79.

79. Sultan, S., et al., D-serine increases adult hippocampal neurogenesis. Front Neurosci, 2013. 7: p. 155.

80. Li, W., X. Xu, and L. Pozzo-Miller, Excitatory synapses are stronger in the hippocampus of Rett syndrome mice due to altered synaptic trafficking of AMPA-type glutamate receptors. Proc Natl Acad Sci U S A, 2016. 113(11): p. E1575–84.

81. Shepherd, J.D. and R.L. Huganir, The cell biology of synaptic plasticity: AMPA receptor trafficking. Annu Rev Cell Dev Biol, 2007. 23: p. 613–43.

82. Goenaga, J., et al., Calcium signaling in astrocytes and gliotransmitter release. Front Synaptic Neurosci, 2023. 15: p. 1138577.

83. Loers, G., et al., Interaction of L1CAM with LC3 Is Required for L1-Dependent Neurite Outgrowth and Neuronal Survival. Int J Mol Sci, 2023. 24(15).

84. Duncan, B.W., K.E. Murphy, and P.F. Maness, Molecular Mechanisms of L1 and NCAM Adhesion Molecules in Synaptic Pruning, Plasticity, and Stabilization. Front Cell Dev Biol, 2021. 9: p. 625340.

85. Navarro-López, J.D., et al., Acquisition-dependent modulation of hippocampal neural cell adhesion molecules by associative motor learning. Front Neuroanat, 2022. 16: p. 1082701.

86. Sytnyk, V., I. Leshchyns’ka, and M. Schachner, Neural Cell Adhesion Molecules of the Immunoglobulin Superfamily Regulate Synapse Formation, Maintenance, and Function. Trends Neurosci, 2017. 40(5): p. 295–308.

87. Shan, L., et al., Astrocyte-Neuron Signaling in Synaptogenesis. Front Cell Dev Biol, 2021. 9: p. 680301.

88. Thompson-Steckel, G. and T.E. Kennedy, Maintaining and modifying connections: roles for axon guidance cues in the mature nervous system. Neuropsychopharmacology, 2014. 39(1): p. 246–7.

89. Mironova, Y.A. and R.J. Giger, Where no synapses go: gatekeepers of circuit remodeling and synaptic strength. Trends Neurosci, 2013. 36(6): p. 363–73.

90. Murai, K.K., et al., Control of hippocampal dendritic spine morphology through ephrin-A3/EphA4 signaling. Nat Neurosci, 2003. 6(2): p. 153–60.

91. Su, Y., et al., Astrocyte endfoot formation controls the termination of oligodendrocyte precursor cell perivascular migration during development. Neuron, 2023. 111(2): p. 190–201.e8.

92. Kıray, H., et al., The multifaceted role of astrocytes in regulating myelination. Exp Neurol, 2016. 283(Pt B): p. 541–9.

93. McEwen, B.S. and S.E. Alves, Estrogen actions in the central nervous system. Endocr Rev, 1999. 20(3): p. 279–307.

94. Finney, C.A., et al., The role of hippocampal estradiol in synaptic plasticity and memory: A systematic review. Front Neuroendocrinol, 2020. 56: p. 100818.

95. Kumar, A., et al., Contribution of estrogen receptor subtypes, ERα, ERβ, and GPER1 in rapid estradiol-mediated enhancement of hippocampal synaptic transmission in mice. Hippocampus, 2015. 25(12): p. 1556–66.

96. Foy, M.R., et al., 17beta-estradiol enhances NMDA receptor-mediated EPSPs and long-term potentiation. J Neurophysiol, 1999. 81(2): p. 925–9.

97. Potier, M., et al., Temporal Memory and Its Enhancement by Estradiol Requires Surface Dynamics of Hippocampal CA1 N-Methyl-D-Aspartate Receptors. Biol Psychiatry, 2016. 79(9): p. 735–745.

98. Boulware, M.I., J.D. Heisler, and K.M. Frick, The memory-enhancing effects of hippocampal estrogen receptor activation involve metabotropic glutamate receptor signaling. J Neurosci, 2013. 33(38): p. 15184–94.

99. Barreto, G., et al., Selective estrogen receptor modulators decrease reactive astrogliosis in the injured brain: effects of aging and prolonged depletion of ovarian hormones. Endocrinology, 2009. 150(11): p. 5010–5.

100. Rummel, J., J.R. Epp, and L.A. Galea, Estradiol does not influence strategy choice but place strategy choice is associated with increased cell proliferation in the hippocampus of female rats. Horm Behav, 2010. 58(4): p. 582–90.

101. Qiu, L.R., et al., Hippocampal volumes differ across the mouse estrous cycle, can change within 24 hours, and associate with cognitive strategies. Neuroimage, 2013. 83: p. 593–8.

102. Warren, S.G., et al., LTP varies across the estrous cycle: enhanced synaptic plasticity in proestrus rats. Brain Res, 1995. 703(1-2): p. 26–30.

