## Supplemental Figure 1 for "Astroglial TNFR2 signaling regulates hippocampal synaptic function and plasticity in a sex dependent manner"

### Supplementary Materials

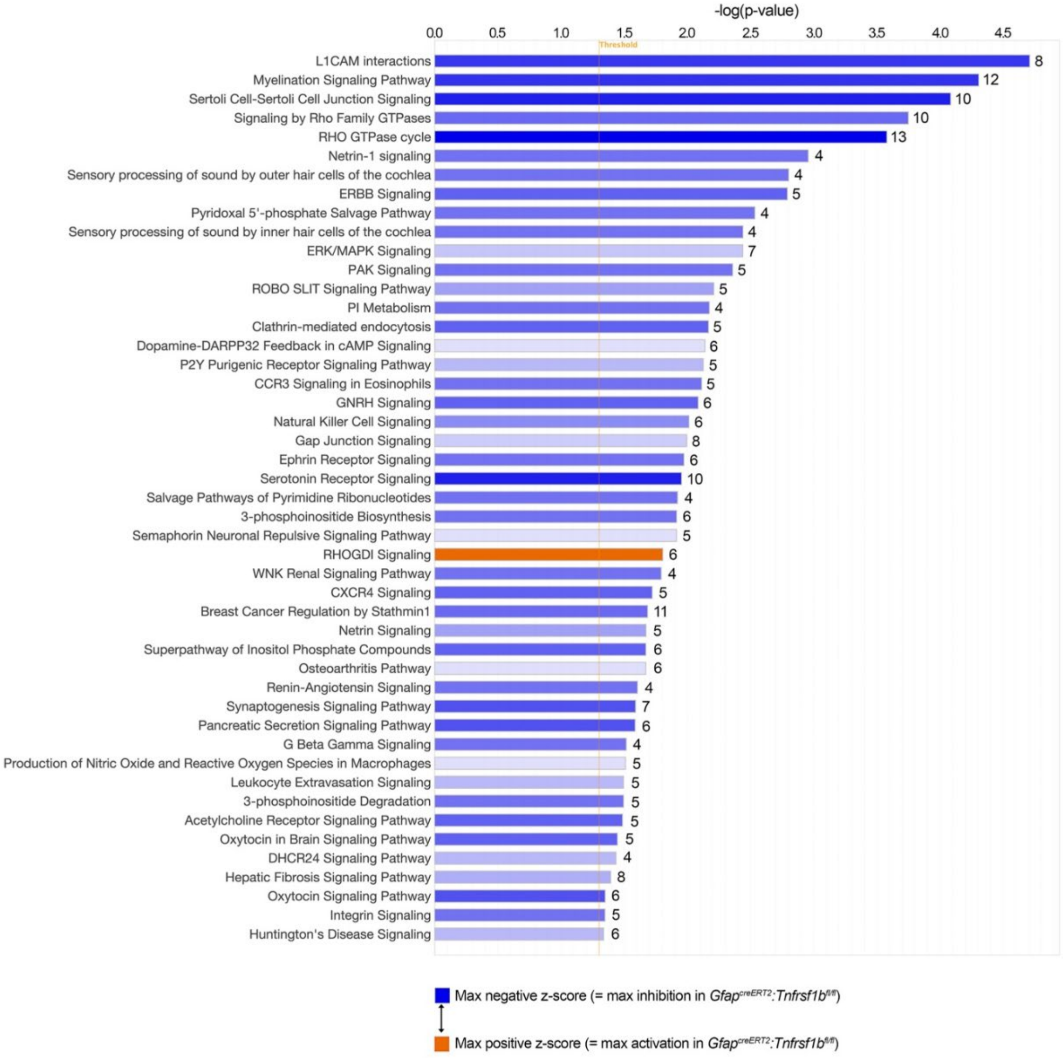


**Supplementary Figure 1. Differentially expressed canonical pathways in *Gfap*$^{creERT2}$*:Tnfrsf1b*$^{fl/fl}$ hippocampal astrocytes.**

Full list of differentially expressed canonical pathways identified by Ingenuity Pathway Analysis (IPA), where genes with an absolute log2 fold change (|LogFc|) ≥ 1.0, and a *p*adj ≤ 0.1 were considered statistically significant. The number of differentially expressed genes in each pathway is depicted on the right of bar graph. Dark blue represents maximum inhibition and dark orange maximum activation in *Gfap*$^{creERT2}$*:Tnfrsf1b*$^{fl/fl}$ hippocampal astrocytes.
